# Identification of the modulatory Ca^2+^ binding sites of acid-sensing ion channel 1a

**DOI:** 10.1101/2023.12.06.570401

**Authors:** Ophélie Molton, Olivier Bignucolo, Stephan Kellenberger

**Author notes:** Corresponding author: Stephan Kellenberger.

## Abstract

Acid-sensing ion channels (ASICs) are neuronal H^+^-gated, Na^+^-permeable channels involved in learning, fear sensing, pain sensation and neurodegeneration. An increase in the extracellular Ca^2+^ concentration shifts the pH dependence of ASIC1a to more acidic values. Here, we predicted candidate residues for Ca^2+^ binding on ASIC1a, based on available structural information and molecular dynamics simulations; the function of channels carrying mutations of these residues was then measured. We identify several residues in cavities previously associated with pH-dependent gating, whose mutation decreased the Ca^2+^-induced shift in ASIC1a pH dependence, likely due to a disruption of Ca^2+^ binding. We show also that Mg^2+^ shares some of the binding sites with Ca^2+^, and that some of the Ca^2+^ binding sites are functionally conserved in the splice variant ASIC1b. Our identification of divalent cation binding sites in ASIC1a shows how Ca^2+^ affects ASIC1a gating, elucidating a regulatory mechanism present in many ion channels.

## Introduction

Acid-sensing ion channels (ASICs) are H^+^-gated and Na^+^-permeable ion channels widely expressed in the nervous system ^1, 2^. They are involved in learning, fear sensing, pain sensation and neurodegeneration after ischemia ^1, 2, 3^. Four ASIC genes encode six different subunits in rodents. Functional ASICs assemble into hetero- and homotrimeric channels whose pH dependence and current kinetics depend on the subunit composition. ASIC1a is the most pH-sensitive subunit in the central nervous system ^1, 2^. The shape of each subunit resembles a hand holding a small ball with the different domains labeled palm, knuckle, finger, thumb and β-ball ^4^ (**Fig. 1a**). The channel contains several electronegative vestibules to which some pharmacological ligands bind, such as the acidic pocket and the central vestibule. Each acidic pocket is enclosed by the thumb, finger and β-ball domains of one subunit and the palm of an adjacent subunit; the central vestibule is located in the palm region ^4, 5, 6^ (**Fig. 1a**).

**Figure 1.**
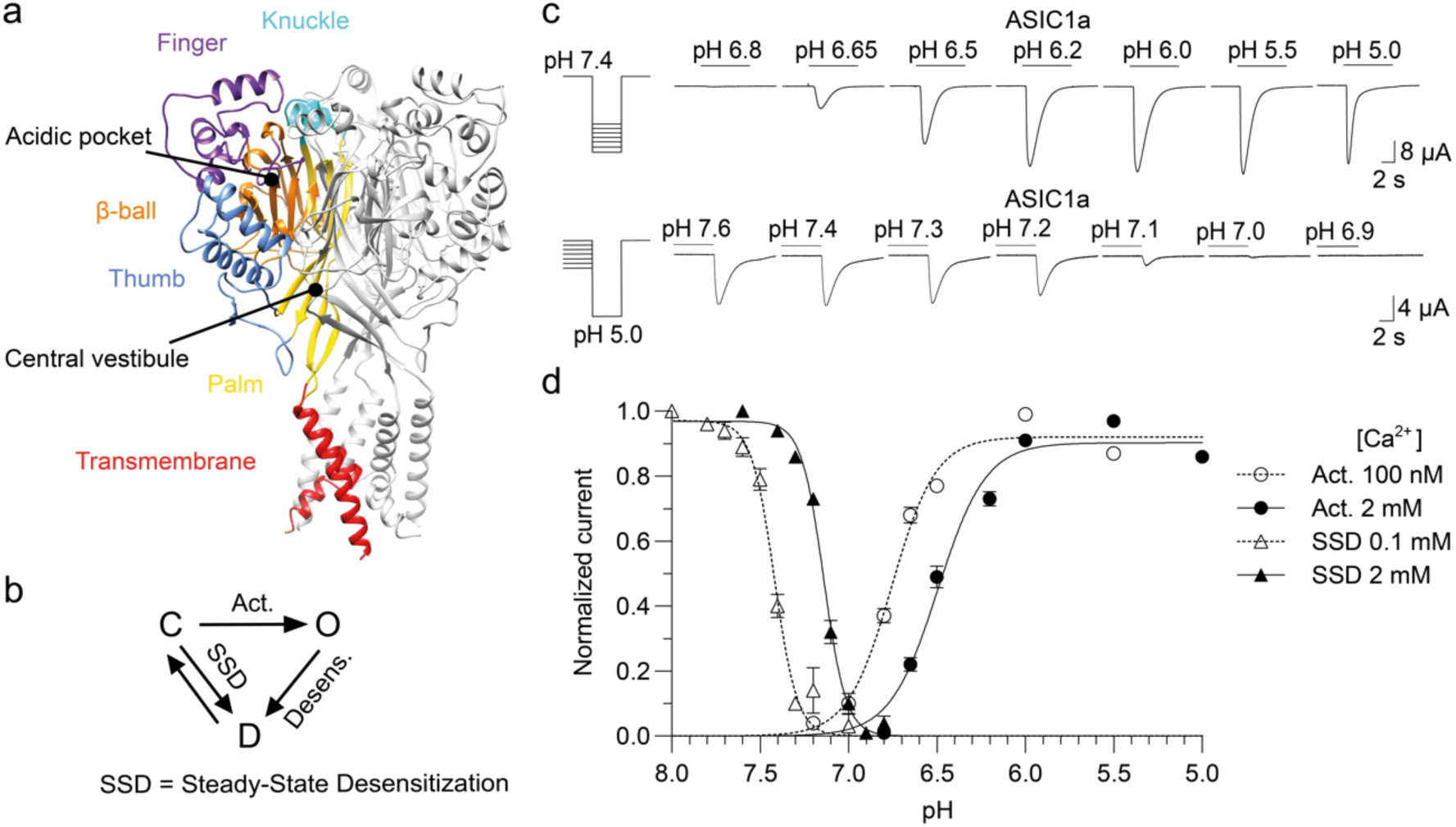
Calcium modulates the pH dependence of activation and steady-state desensitization (SSD) of ASIC1a. **a** Structural image of ASIC1a trimer, based on a model of the crystal structure (PDB number 5WKU). The different domains are indicated by specific colors in one of the three ASIC subunits. **b** Kinetic scheme of ASIC1a with the three functional states (closed (C), open (O) and desensitized (D)) and the transitions corresponding to activation, desensitization from the open state and SSD indicated by arrows. **c** Representative current traces of *Xenopus laevis* oocytes expressing ASIC1a WT were obtained by two-electrode voltage clamp to -60 mV to determine the pH dependence of activation (upper panel) and the pH dependence of SSD (lower panel). Channels were activated every 60s by a 10-s perfusion with stimulation solution. For the measurement of the activation pH dependence, stimulation solutions of varying pH were applied, and the pH of the conditioning solution, perfused between stimulations was 7.4. For the measurement of the SSD pH dependence, solutions of varying pH were applied for 50 s before the stimulation by pH5.0. **d** pH dependence curves of activation and SSD. Activation curves were measured at 100 nM free Ca^2+^ (open circles) or 2 mM Ca^2+^ in the stimulation solution (filled circles) with a conditioning solution containing 2 mM Ca^2+^. SSD curves were measured at 0.1 mM Ca^2+^ (open triangles) or 2 mM Ca^2+^ in the conditioning solution (filled triangles), with 2 mM Ca^2+^ in the stimulation solution. Currents were normalized to the maximum peak current induced. The lines represent fits to the Hill equation (n = 17-56).

Extracellular free Ca^2+^ concentrations decrease locally during high neuronal activity, during seizures and in ischemic stroke ^7, 8, 9^. Lowering of the Ca^2+^ concentration affects the activity of many neuronal ion channels ^10, 11, 12^ and thereby neuronal excitability ^13^. Exposure to an acidic extracellular pH leads to a transient opening of ASICs, followed by a current decay due to the entry into a non-conducting desensitized state (**Fig. 1b-c**) ^14^. Calcium ions appear to compete with protons for binding sites in ASICs, since an increase in their concentration shifts the pH dependence towards more acidic pH values ^15, 16^. In addition to its effects on gating, extracellular Ca^2+^ also inhibits ASIC1a currents by a pore block by binding into the ion pore, with an IC_50_ of the order of millimolar ^17^. In ASIC3, two different Ca^2+^ binding sites are involved in channel regulation, one in the pore and one in the acidic pocket ^18, 19^. For ASIC1a, the understanding of gating modulation by Ca^2+^ remains incomplete because the Ca^2+^ binding sites mediating the shift in pH dependence in ASIC1a have not been identified yet.

A recent study showed approximate locations of Ca^2+^ binding in the acidic pocket and central vestibule of chicken ASIC1a, based on crystals that were soaked in Ba^2+ 20^. These two cavities had been shown to contain numerous proton binding sites ^21, 22, 23, 24, 25^ and to undergo substantial conformational changes during gating ^6, 26, 27, 28, 29, 30^. In the high pH resting state structure, two divalent ions bind to each acidic pocket, while three divalent ions bind to the central vestibule. In the low pH desensitized state, the divalent cation binding sites in the central vestibule are lost and the number of binding sites in each acidic pocket is reduced from two to one.

Here we have carried out molecular dynamics (MD) simulations to refine the Ca^2+^ coordination in an ASIC1a structural model, identifying, out of the many negatively charged residues, specific residues in the acidic pocket and the central vestibule as the most likely binding sites of Ca^2+^. We mutated candidate residues and measured the modulation of the pH dependence by extracellular Ca^2+^. We identified several residues in the acidic pocket, the central vestibule and the pore entry that contribute to the modulatory effect of Ca^2+^ and are most likely part of binding sites. In addition, we show that Mg^2+^ shares binding sites with Ca^2+^ for desensitization, and that Ca^2+^ binding sites for desensitization in the central vestibule are functionally conserved between the splice variants ASIC1a and ASIC1b.

## Results

### Calcium modulates ASIC1a function

To investigate the effects of Ca^2+^ on ASIC1a activity, the pH dependence of activation and steady-state desensitization (SSD) was measured at two different extracellular Ca^2+^ concentrations. Wild-type (WT) or mutant ASIC1a channels were expressed in *Xenopus laevis* oocytes, and their function was measured by two-electrode voltage-clamp. The pH dependence of activation (“Act.” in **Fig. 1b**) was measured by stimulating the channels once per minute during 10 s to a series of increasingly acidic pH solutions, as illustrated in the top panel of **Fig. 1c**. The conditioning pH between stimulations was 7.4 in all experiments unless noted. The Ca^2+^ concentration was kept at 2 mM in the conditioning solution, while it was either 2 mM or 100 nM in the stimulation solution. In **Fig. 1d**, normalized currents measured with this protocol are plotted as spheres as a function of the stimulation pH. Decreasing the extracellular Ca^2+^ concentration shifted the pH dependence to more alkaline values. Fitting the pH dependence curves yielded pH values of half-maximal activation (pH_50_) of 6.50 ± 0.12 (n = 50) at 2 mM Ca^2+^ and 6.75 ± 0.11 (n = 53) at 100 nM Ca^2+^, indicating a shift toward more alkaline pH_50_ values by 0.25 pH units with the lower Ca^2+^ concentration. The pH dependence of steady-state desensitization (SSD), the transition from the closed to the desensitized state without apparent opening (**Fig. 1b**) was measured by a 10-s stimulation by pH5.0 once per minute, which was preceded by 50-s exposures to conditioning solutions of increasingly acidic pH (**Fig. 1c**, lower panel). This protocol measures the channel availability for activation after exposure to the indicated conditioning pH. The Ca^2+^ concentration was either 2 mM or 0.1 mM in the conditioning solutions and was kept constant at 2 mM in the stimulation solution. Normalized current amplitudes are plotted as triangles as a function of the conditioning pH in **Fig. 1d**. The pH of half-maximum desensitization (pHD_50_) was 7.12 ± 0.04 (n = 22) at 2 mM Ca^2+^ and 7.39 ± 0.06 (n = 23) at 0.1 mM Ca^2+^, indicating a shift toward more alkaline pHD_50_ value by 0.27 pH units with the lower Ca^2+^ concentration. Thus, lowering the Ca^2+^ concentration shifts the ASIC1a pH dependence of activation and SSD to more alkaline values, as previously shown ^15^.

### Prediction of candidates for Ca^2+^ coordination by MD simulations

The two-fold positively charged Ca^2+^ ions tend to bind to negatively charged residues which contain carboxylate groups. To obtain a more precise prediction of divalent-coordinating residues than provided from the structure of the Ba^2+^-soaked crystals ^20^ and identify residues participating in the coordination of divalent cations, MD simulations were carried out with a human ASIC1a model of the chicken ASIC1a resting structure (PDB code 6VTL) ^26^, in which two Ca^2+^ residues per acidic pocket, and a total of three residues in the central vestibule were placed according to their approximate published location ^20^. The protonation state of the titratable side chains was updated every 100 ns to mimic pH7.4, as described (*Methods*) during the MD simulations which were carried out for 600 ns using a system containing two independent channels. During these simulations, the proximity of the Ca^2+^ ions to acidic residues (Asp, Glu) was monitored and the residues interacting more than 10% of each period of 100 ns were included in a more detailed analysis (*Methods*).

Sixteen residues per subunit fulfilled this criterium, 12 in the acidic pocket and 4 in the central vestibule. Of these residues, Ca^2+^ - side chain distances (distance between the center of the Ca^2+^ atom and the gravity center of the two oxygen residues of the carbonyl group) were measured every 400 ps. **Fig. 2a** plots for each residue the fraction of time at which the distance to the closest Ca^2+^ ion was < 4 or < 6 Å, respectively. **Fig. 2b** shows the mean distance of a given residue to the closest Ca^2+^ ion. For this analysis, only Ca^2+^ - side chain distances ≤ 10 Å were considered. Residues that had during ≥ 50% of the simulation time a Ca^2+^ ion in their proximity, displayed relatively short (∼5 Å) distances to the closest Ca^2+^ ion. This analysis predicted E97, E219, E238, E242, D347, D351 and D409 of the acidic pocket and E375, E413 and E418 of the central vestibule as good candidates for Ca^2+^ binding and D237 and E355 (acidic pocket) and E79 (central vestibule) as lower priority candidates (**Fig. 2a-b**). The location in the structure of these residues is shown in **Figs. 2c-d**. Residues facing the external part of the protein or being distant from other acidic residues, such as E177, showed lower Ca^2+^ interactions in the MD simulations and were therefore not functionally investigated.

**Figure 2:**
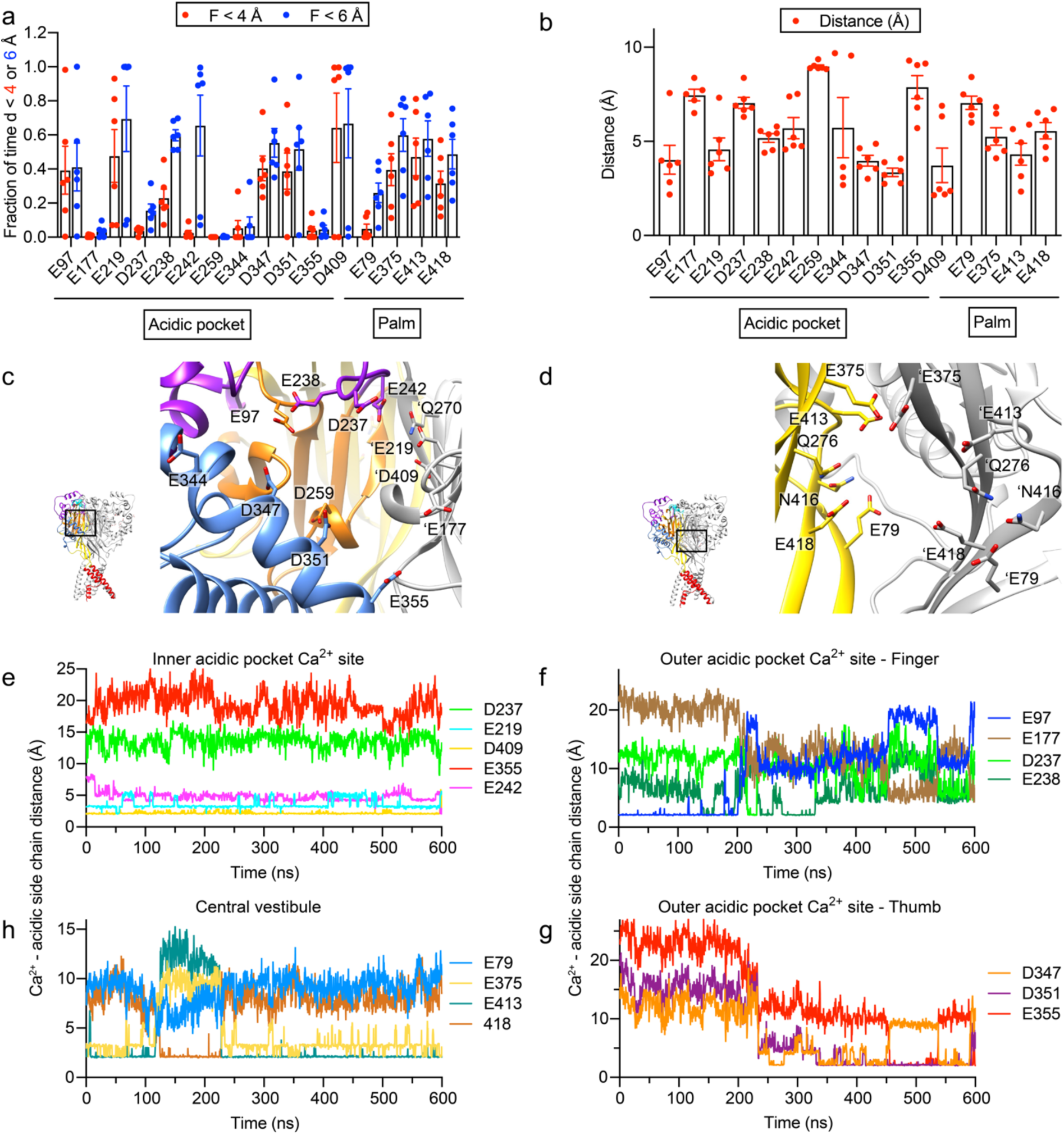
MD simulations predict Ca^2+^-binding residues. MD simulations were carried out over a total duration of 600 ns with two independent ASIC1a channels, containing thus 6 acidic pockets with 2 Ca^2+^ ions each, and 6 subunit-Ca^2+^ pairs in the central vestibule (*Materials and Methods*). If distances, measured between Ca^2+^ ions and the center-of-mass of the carbonyl groups of acidic side chains during a 100-ns stretch were during ≥ 10% of the time < 10 Å, they were considered for the analysis, carried out every 400 ps. **a** Fraction of time during which a Ca^2+^ ion was closer than 4 Å (red symbols) or 6 Å (blue symbols) to the indicated residue. **b** Distance between Ca^2+^ ions and the center-of-mass of the carbonyl groups of acidic side chains. For this analysis, distances >10 Å were excluded. **a-b** each data point represents the measurement of a different acidic pocket, or a different central vestibule subunit-Ca^2+^ ion pair. **c-d** Structural images of the acidic pocket **c** and the central vestibule **d** in the closed conformation (model based on PDB 5WKU), showing the residues for which distances to Ca^2+^ ions were measured in **a-b**. Additional residues, not included in the MD analysis, but studied by functional experiments, are shown, Q270 (close to E219 and E242; **c**) and Q276 and N416 (close to E413 and E418 for Q276 and E79 for N416, **d**). Amino acid residue labels without prefix and labels with the prefix ’ are on different subunits. In **d**, only two subunits are shown. (**e-h**), Time course of Ca^2+^-side-chain distances for two Ca^2+^ ions in the same acidic pocket, one placed at the inner AP Ca^2+^ site **e**, one placed at the outer AP Ca^2+^ site (**f-g**, with distances to finger (**f**) and thumb residues (**g**)), and of a Ca^2+^ ion placed in the central vestibule (**h**). These are representative traces out of 6 measurements each in the acidic pocket and for Ca^2+^ ion central vestibule subunit pairs.

Besides guiding the choice of residues of interest for the functional analysis, the MD simulations provided information on the dynamics of Ca^2+^ binding in both the acidic pocket and the central vestibule. Since the simulations were carried out in two separate ASIC channels, they provided information on the dynamics of six acidic pockets with two Ca^2+^ ions each, and six subunit-Ca^2+^ ion pairs of the central vestibule. In the acidic pocket, the Ca^2+^ ion placed at the beginning of the simulation close to E219 and D409, which we name here the “inner AP Ca^2+^ site” stayed in 4/6 simulations throughout the entire simulation in close proximity of E219, D409 and generally, at an increased distance, to E242, as illustrated for a typical experiment in **Fig. 2e**. In the two other simulations, the Ca^2+^ moved towards residues D351 and D347, and left the acidic pocket after about 300 ns. The second Ca^2+^ ion was placed in proximity of E97, named here “outer AP Ca^2+^ site”. In 5/6 simulations it moved during the simulation towards the center of the acidic pocket into proximity of D347 and D351, or E238, where it stayed for the rest of the simulation (**Fig. 2f-g**). In one simulation, the Ca^2+^ ion stayed throughout the simulation close to E97.

In the central vestibule, the Ca^2+^ ion close to a given subunit changed in two simulations frequently positions relative to E79, E375, E413 and E418, without showing any preference for a given residue. In two simulations, it stayed for most of the time close to E375 and E413, as illustrated in **Fig. 2h**. In two simulations, the Ca^2+^ ion stayed for ∼400 ns close to E375 and E413 and in part to E418, before switching its position with the Ca^2+^ ion initially positioned close to the adjacent subunit (**Supplementary Fig. 1**).

### Mutations in the acidic pocket and the central vestibule decrease the effect of Ca^2+^ on ASIC1a activation

Thirteen negatively charged residues were selected as candidate residues for Ca^2+^ binding sites of ASIC1a based on the MD simulations and their position in the ASIC1a structure. These residues were mutated individually to Ala. In addition, two Gln and one Asn were also included as candidates for Ca^2+^-binding sites, since they are in proximity of some candidates identified by MD simulations and could interact with Ca^2+^ ions with their partial negative charge (**Fig. 2c**-**d**). Alterations of the charge and size of side chains by the mutation to Ala should decrease the ability of Ca^2+^ to bind and compromise its effect on the ASIC1a pH dependence in case the mutated residue is part of a Ca^2+^-binding site. The pH dependence was determined in the same oocyte at two Ca^2+^ concentrations. At 2 mM Ca^2+^, the pH_50_ values of many mutants were significantly lower than the WT values (**Fig. 3a-b**), indicating a decreased pH sensitivity. The ASIC1a WT ΔpH_50_, measured by subtracting the pH_50_ at 2 mM from the one measured at 100 nM Ca^2+^, was 0.25 ± 0.09 (n = 50; **Fig. 3c**). In the acidic pocket, the mutations E238A and D347A decreased the ΔpH_50_ significantly by 41% and 48% respectively, compared to the WT (**Fig. 3c**). The mutations E219A and Q270A showed a tendency towards smaller ΛλpH_50_ that was however not statistically significant. A channel construct containing these four mutations, termed “AP-Act.”, showed a significant decrease of the ΔpH_50_ compared to the WT (0.10 ± 0.09, n = 8; **Fig. 3d**), although the shift in pH_50_ was not completely suppressed. The mutant D409A showed a significant increase of the ΔpH_50_ compared to the WT (0.41 ± 0.09, n = 10), suggesting that this residue does not favor Ca^2+^ coordination or the competition with protons in the context of activation (**Fig. 3c**). In the central vestibule, the mutants E79A and E418A showed the strongest deviation from WT, displaying no significant modulation of the pH_50_ by Ca^2+^ (**Fig. 3a** and **3c**). In addition, the mutations Q276A, E375A and E413A decreased the ΔpH_50_ significantly by 38-63% (**Fig. 3c**). These effects were significantly smaller than the decrease in ΔpH_50_ induced by the mutation E79A (Q276A, p < 0.01; E375A, p < 0.0001; E413A, p < 0.001; ANOVA and Tukey’s multiple comparisons test).

**Figure 3.**
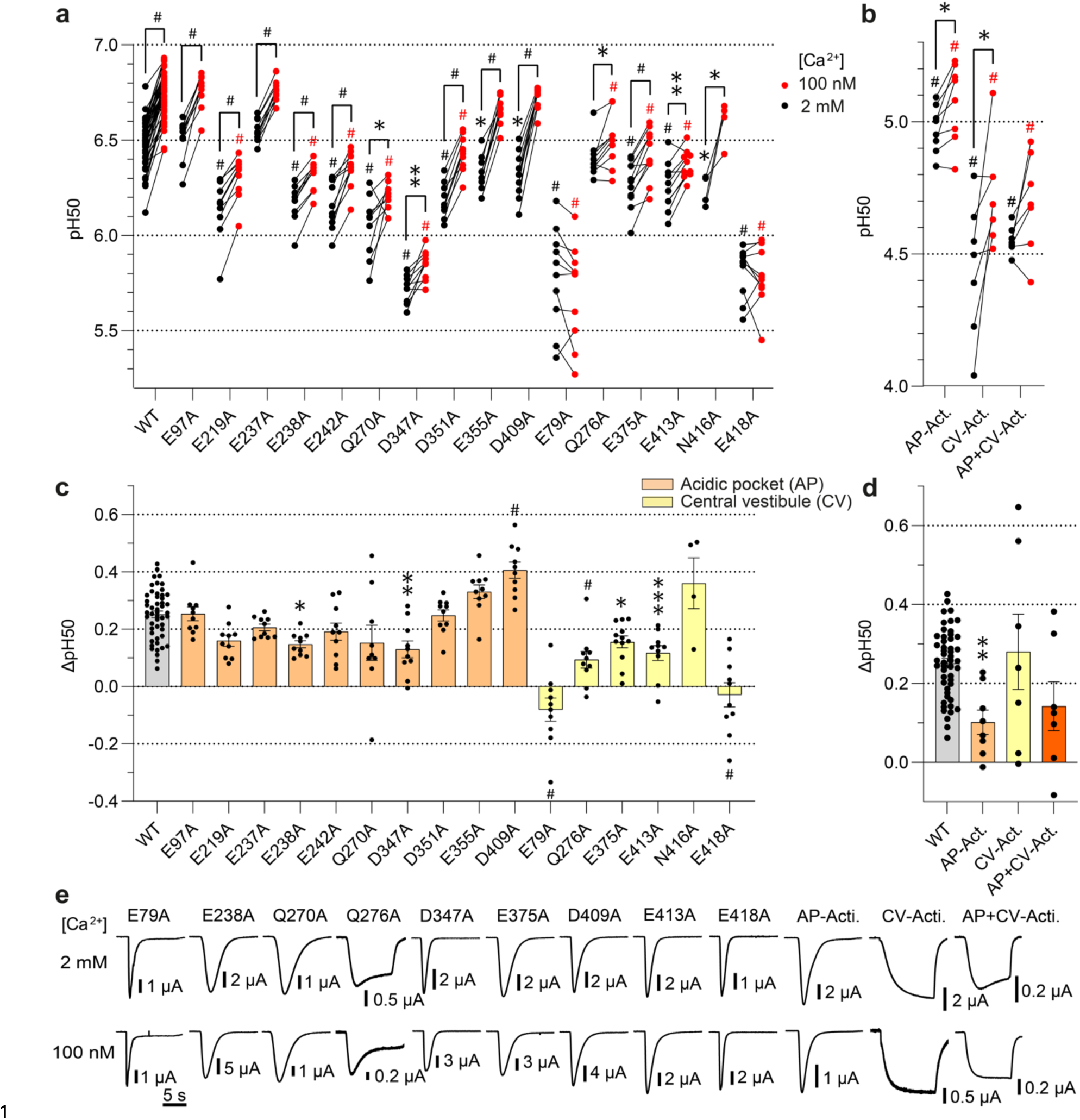
Functional analysis of mutants identifies Ca^2+^ binding sites for activation in the acidic pocket and the central vestibule. **a-b** pH values for half-maximal activation (pH_50_) were obtained from activation curves with stimulation solutions containing 100 nM free Ca^2+^ (red symbols) or 2 mM Ca^2+^ (black symbols), n = 4-53. The pH dependence at 100 nM and 2 mM free Ca^2+^ was measured in the same oocytes. The mutant AP-Act. contains the mutations E219A, E238A, Q270A and D347A. CV-Act. contains the mutations E79A, Q276A, E375A, E413A, and E418A. The comparison between pH_50_ at 100 nM free Ca^2+^ and 2 mM Ca^2+^ was done by paired t-test. For comparisons of pH_50_ values of mutants relative to the WT at 100 nM or 2 mM Ca^2+^, a one-way ANOVA and Tukey’s multiple comparison test were carried out. *, p < 0.05; **, p < 0.01; ***, p < 0.001; #, p < 0.0001. **c-d** ΔpH_50_ (pH_50,100nM_-pH_50,2mM_) values are shown as mean ± SEM, n = 4-50. Comparison of the mutants to the WT was done by one-way ANOVA and Dunnett’s multiple comparison test. *, p < 0.05; **, p < 0.01; ***, p < 0.001; #, p < 0.0001. **e** Representative current traces of mutants showing a significant reduction in the ΔpH_50_ relative to the WT and of the combined mutants. The traces were obtained at the two Ca^2+^ concentrations, at a pH close to the pH_50_.

A channel containing these 5 mutations of the central vestibule that individually had produced a significant reduction of the ΔpH_50_, “CV-Act.”, had only a sustained current (traces in **Fig. 3e**), although individual mutations except for Q276A and E79A had not induced any sustained currents. In addition, the pH dependence was strongly shifted towards acidic values (**Fig. 3b**). The CV-Act. mutant showed no significant difference of the Ca^2+^-dependent ΔpH_50_ compared with the WT (0.28 ± 0.25, n = 7; **Fig. 3d**). On a construct combining the four acidic pocket and five central vestibule mutations, termed “AP+CV-Act.” the modulation by Ca^2+^ was variable and not significantly different from WT (ΔpH_50_ = 0.18 ± 0.17, n = 10). This mutant also produced a completely sustained current (**Fig. 3e**). We had previously observed sustained currents and strongly shifted pH dependence when mutations of the palm/central vestibule were combined ^27^, suggesting that these combined mutants open to an alternative open state. Since the basic current properties of the CV-Act. and AP+CV-Act. mutants are profoundly different from the WT, they cannot be used to infer how the combination of mutations would affect the default opening process of ASIC1a. Although the combination of central vestibule mutations was not conclusive, the analysis of the activation shows that two mutations in the acidic pocket and five mutations in the central vestibule decrease the Ca^2+^-dependent ΔpH_50_ compared to the WT.

### Acidic pocket and central vestibule mutations decrease the Ca^2+^ modulation of SSD

The pHD_50_ values of SSD of many mutants were also significantly different from WT (**Fig. 4a**-**b**). The WT ΔpHD_50_ was 0.28 ± 0.03 (n = 17, **Fig. 4c**). A significantly decreased ΔpHD_50_ compared to WT was observed for the seven acidic pocket mutants, E97A, E219A, E238A, E242A, Q270A, D347A and D409A (**Fig. 4c**). The ΔpHD_50_ was decreased by these mutations by 14-32%. In the central vestibule, only the mutants E375A and E413A decreased the ΔpHD_50_ values significantly, by 22% and 24%, respectively (**Fig. 4c**). The combination of the seven mutations in the acidic pocket or of the two mutations in the central vestibule that had produced a significant reduction of the ΔpHD_50_, termed “AP-SSD” and “CV-SSD”, showed a significant decrease of the ΔpHD_50_ compared to the WT to 0.05 ± 0.1 (n = 8) and 0.13 ± 0.03 (n = 8) respectively (**Fig. 4d**). These two combined mutants produced transient currents (**Fig. 4e**) and showed a greater reduction of the ΔpHD_50_ in comparison to the individual mutants (ANOVA and Dunnett’s multiple comparisons test; AP-SSD compared with all individual acidic pocket mutants, p < 0.0001; Palm-SSD compared with E375A and E413A, p < 0.0001). The construct combining the mutations of AP-SSD and CV-SSD was non-functional, which precluded the investigation of the effect of combining all the best candidates.

**Figure 4.**
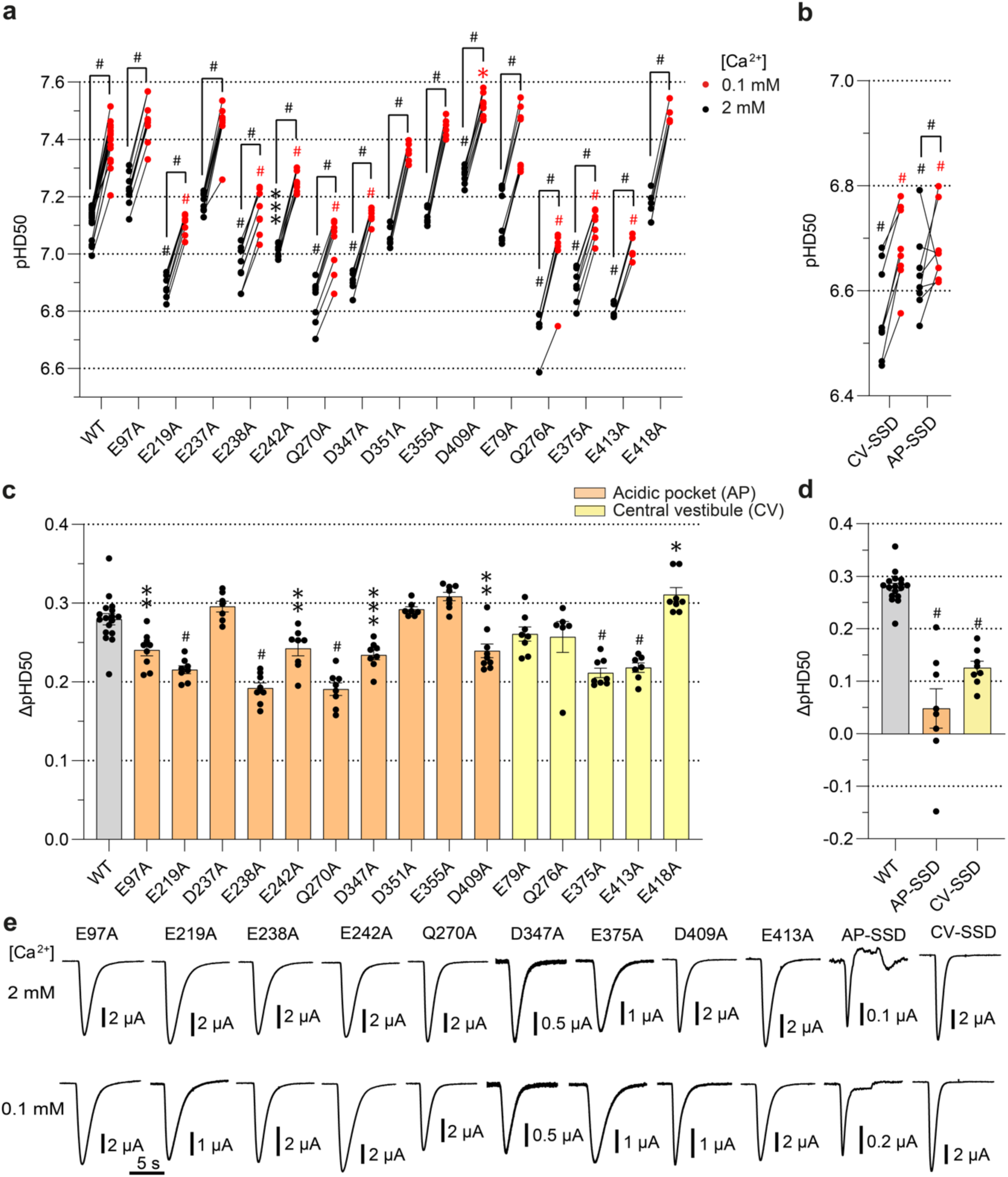
Mutations in the acidic pocket and the central vestibule affect the modulation of the SSD by Ca^2+^. **a-b** pH values for half-maximal SSD (pHD_50_) obtained from SSD curves with conditioning solutions containing 0.1 mM Ca^2+^ (red symbols) or 2 mM Ca^2+^ (black symbols), n = 6-17. The pH dependence at 0.1 mM and 2 mM Ca^2+^ was measured in the same oocytes. AP-SSD combines the mutations E97A, E219A, E238A, Q270A, E242A, D347A and D409A. CV-SSD combines the mutations E375A and E413A. **c-d** ΔpHD_50_ (pHD_50,0.1mM_-pHD_50,2mM_) values are shown as mean ± SEM, n = 6-17. **e** Representative current traces of mutants showing a significant reduction in the ΔpHD_50_ relative to the WT. The traces were obtained at the two free Ca^2+^ concentrations, at a pH close to the pHD_50_. **a-d**, The same statistical tests as in Fig. 3 were carried out.

### Known Ca^2+^ binding sites in the pore entry are also involved in the Ca^2+^ modulation of the ASIC1a pH dependence

In addition to modulating the pH dependence, Ca^2+^ has been shown to inhibit ASIC1a by a pore block. Two residues in the pore entry of rat ASIC1a, E425 and D432, were shown to contribute to this effect ^17^. We tested whether these two residues were involved as well in the modulation of the pH dependence by Ca^2+^. The corresponding residues in human ASIC1a, E427 and D434 (**Fig. 5a**), were mutated to Ala. First, the inhibition by Ca^2+^ was measured by exposing WT and mutants to increasing extracellular Ca^2+^ concentrations at pH5.5, at which the channels are fully activated. In ASIC1a WT, 10 mM Ca^2+^ inhibited 51 ± 3% of the maximum current amplitude (**Fig. 5b-c**). The inhibition by 10 mM Ca^2+^ was 35 ± 2% with E427A and 16 ± 10% with D434A. For D434A, this reduction was significantly different from the WT (p < 0.001), thus similar to the results obtained with the rat ASIC1a mutants ^17^. Next, the involvement of these residues in the modulation of the pH dependence by Ca^2+^ was assessed. At 2 mM Ca^2+^, the pH_50_ and pHD_50_ of the two mutants were very similar to the corresponding WT values (**Fig. 5d-e**). The mutation E434A decreased the ΔpH_50_ of activation compared to the WT by about half (**Fig. 5d**). The mutation E427A did not affect Ca^2+^ modulation of activation; it reduced however the ΔpHD_50_ as compared to WT by 23% (**Fig. 5e**). This shows that in addition to the Ca^2+^ pore block, the residues E427 and D434 are involved in modulating the pH dependence of human ASIC1a by Ca^2+^.

**Figure 5.**
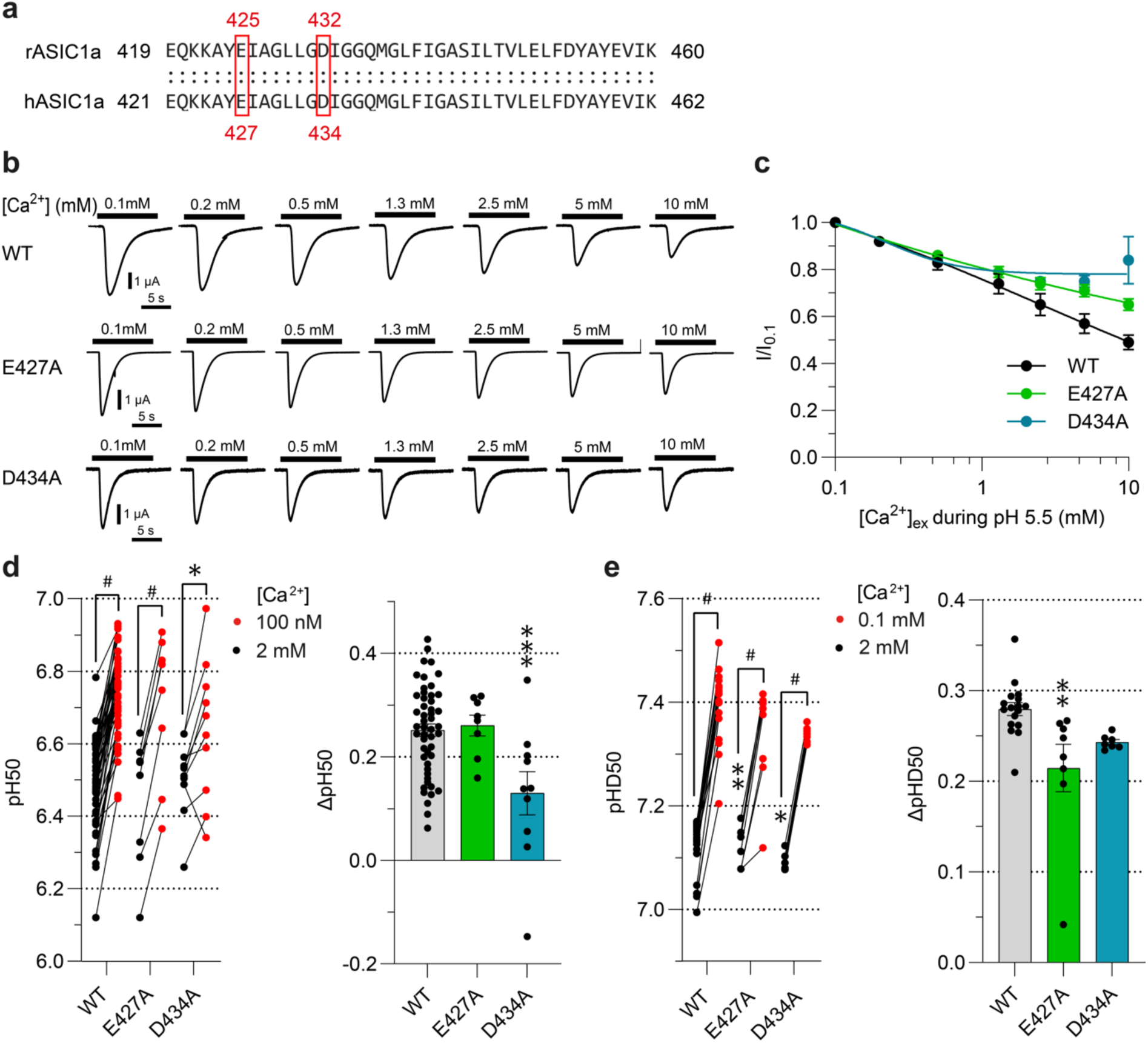
Mutations of two amino acid residues in the pore reduce Ca^2+^ block and modulation of pH dependence by Ca^2+^. **a** Alignment of amino acid sequences of rat ASIC1a and human ASIC1a at the level of the transmembrane domain TM2. Corresponding conserved residues are highlighted by red frames. **b** Representative current traces of ASIC1a WT and mutants obtained by application of increasing Ca^2+^ concentrations during 10 s stimulation solutions at pH5.5, as indicated. The Ca^2+^ concentration in the conditioning solution of pH7.4 was 2mM. **c** Concentration-response curves for Ca^2+^ inhibition of ASIC1a WT and mutants. The lines represent fits to an inhibition equation. **d-e** Left panels, pH_50_ and pHD_50_ values obtained from activation and SSD curves with stimulation (**d**) or conditioning solutions (**e**) containing low free Ca^2+^ as indicated (red symbols) or 2 mM Ca^2+^ (black symbols), n = 7-53. The pH dependence at the two Ca^2+^ concentrations was measured in the same oocytes. A paired t-test was used to compare the pH_50_ or pHD_50_ values between the two Ca^2+^ conditions, and a one-way ANOVA with Tukey’s multiple comparison test to compare the pH_50_ or pHD_50_ values of mutants to the WT in the corresponding Ca^2+^ condition. Right panels, ΔpH_50_ (pH_50,100nM_-pH_50,2mM_, **d**) or ΔpHD_50_ (pHD_50,0.1mM_-pHD_50,2mM,_ **e**) are shown. Comparison of the mutants to the WT was done by one-way ANOVA and Dunnett’s multiple comparison test. *, p < 0.05; **, p < 0.01; ***, p < 0.001; #, p < 0.0001.

### Stabilization of the ASIC1a resting state by Ca^2+^ demonstrated by mutations of two central vestibule residues

Increasing the extracellular Ca^2+^ concentration from 0.1 mM to 2 mM shifted the pH dependence of SSD to more acidic values for ASIC1a (**Fig. 1d**), thereby increasing the number of available channels at a given pH. It is expected that an increase in Ca^2+^ concentration will, at a given pH, increase the rate of recovery and/or slow the rate of desensitization from the closed state ^15^. The central vestibule is enclosed by the lower palm domain of the three subunits, which has been shown to be involved in desensitization from open and closed states ^31^. We have shown that the residue E375 and E413, located in the central vestibule, are involved in Ca^2+^ coordination modulating SSD (**Fig. 4c**). To further investigate their roles in the stabilization of the ASIC1a resting state by Ca^2+^, the rate of recovery from desensitization after exposure to pH6.0 was measured by exposing the desensitized channels during increasing periods to conditioning pH7.4 with a Ca^2+^concentration of 2 mM or 0.1 mM, before a second stimulation by pH6.0 (**Fig. 6a**). For WT ASIC1a, a slower time constant of recovery from desensitization was observed at 0.1 mM Ca^2+^ (22.8 ± 4.9 s, n = 10) compared with 2 mM Ca^2+^ (4.0 ± 1.0 s, n=10; **Fig. 6b**). As a measure of the modulatory effect of Ca^2+^, the τ_0.1mM Ca_/τ_2mM Ca_ ratio was calculated in each experiment, which was 8.3 ± 2.0 for the WT (**Fig. 6c**, n = 10). The pH during the recovery period of ASIC1a WT was 7.4 with both Ca^2+^ conditions, which is 0.28 pH units above its pHD_50_ in the presence of 2 mM Ca^2+^ (**Figs. 4a** and **1d**). Since the pH dependence of SSD of the two mutants was shifted to more acidic values, the pH in the conditioning solutions was adapted for the mutants, to pH7.1 for E375A and pH7.0 for E413A. At 2 mM Ca^2+^, E413A showed a time constant of recovery that was very close to that of the WT, while the recovery kinetics of E375A were with 8.9 ± 2.5 s (n = 7) somewhat slower. The effect of the Ca^2+^ concentration change was significantly smaller in the mutants, with a τ_0.1mM Ca_/τ_2mM Ca_ ratio of 1.5 ± 0.2 (n = 7) for E375A and 1.8 ± 0.2 (n = 8) for E413A (**Fig. 6c**), as further illustrated by smaller shifts in the recovery curves (**Fig. 6b**).

**Figure 6.**
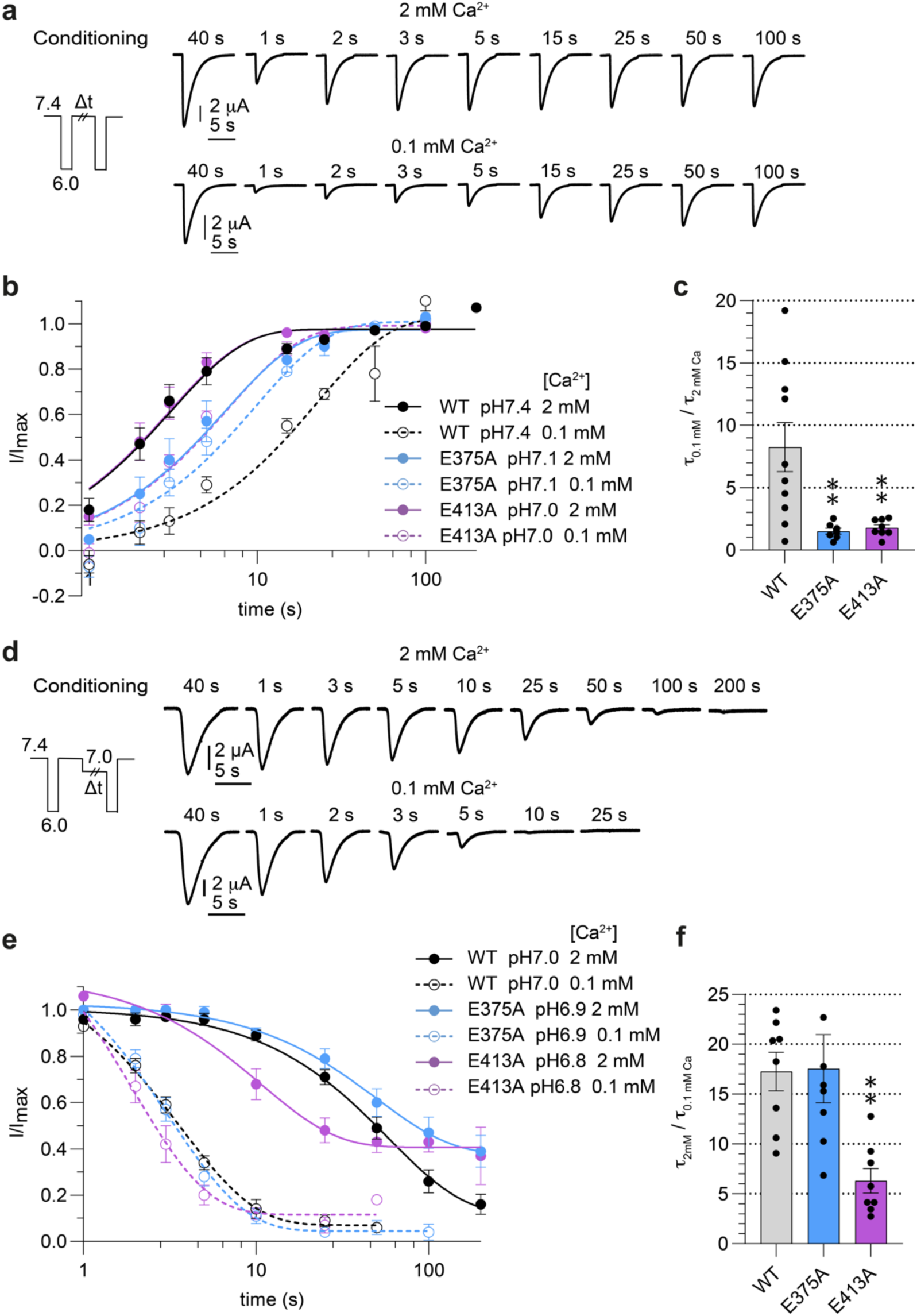
Mutations affecting Ca^2+^ modulation change the kinetics of transitions in and out of the desensitized state. **a** Representative current traces of ASIC1a WT recovery from desensitization experiments, following the protocol illustrated on the left. Two stimulations to pH6.0 were separated by an interval of increasing duration at the conditioning solution of pH7.4 containing 2 mM Ca^2+^ (top) or 0.1 mM Ca^2+^ (bottom panel). The duration of the conditioning interval at pH7.4 is indicated above each trace. **b** Current peak amplitudes normalized to the control current amplitude of each recovery protocol are plotted as a function of the duration at the conditioning solution between the two paired stimulations to pH6.0, n = 7-10. The connecting lines are fits to a single exponential. The conditioning pH used for the different constructs is indicated in the figure. **c** τ ratios (τ_0.1mM Ca_ / τ_2mM Ca_) calculated from the fits to the recovery time course, n = 7-10. Comparison between the mutants and the WT was done by one-way ANOVA and Dunnett’s multiple comparison test. *, p < 0.05; **, p < 0.01; ***, p < 0.001; #, p < 0.0001. **d** Representative ASIC1a current traces from the onset of SSD protocol, as illustrated on the left. A 3.5-s pH6.0 stimulation (control response) was applied before the application of pH7.4 for 50 s to allow the channel to recover completely from desensitization. Subsequently, the WT channels were exposed to a conditioning pH7.0 whose duration was increased in each round, followed by a second stimulation with pH6.0. The test conditioning solution contained 2 mM (top) or 0.1 mM Ca^2+^ (bottom panel). The duration of the conditioning period is indicated above each trace. **e** Current peak amplitudes normalized to the control current amplitude of each desensitization onset protocol, plotted as a function of the conditioning period, n = 8. The conditioning pH used for the different constructs is indicated in the figure. The connecting lines represent exponential fits. **f** τ ratios (1_2mM Ca_ / τ_0.1mM Ca_) obtained from the fit of the normalized current peak amplitudes of the onset of SSD protocol, n = 8. Comparison of the mutants to the WT was done by one-way ANOVA and Dunnett’s multiple comparison test. *, p < 0.05; **, p < 0.01; ***, p < 0.001; #, p < 0.0001.

To study the inverse transition, the onset of SSD, channels were exposed during varying periods to a pH that induces desensitization, pH7.0 for WT, pH6.9 for E375A and pH6.8 for E413A (**Fig. 6d**). These pH values were chosen because they desensitize the channels by ∼50% after a 50-s exposure to the conditioning pH at 2 mM Ca^2+^. The fraction of channels having entered the desensitized state was determined as the ratio of the pH6.0-induced current measured after exposure to the desensitizing pH / control current amplitude (**Fig. 6e**). In WT ASIC1a, the onset of SSD at 0.1 mM Ca^2+^ (1_0.1mM Ca_ = 3.9 ± 0.4 s, n = 8) was faster than in the presence of 2 mM Ca^2+^ (1_2mM Ca_ = 63.6 ± 8.0, **Fig. 6e-f**) with a τ_2mM Ca_/τ_0.1mM Ca_ ratio of 17.3 ± 1.9 (n = 8). This is expected since lowering of the Ca^2+^ concentration shifts the pHD_50_ to more alkaline values. Ca^2+^ induced a similar shift as in WT also in E375A, while in E413A, the τ_2mM Ca_/τ_0.1mM Ca_ ratio was with 6.3 ± 1.2 (n = 8) significantly lower than the WT value. In conclusion, both mutations impair the Ca^2+^ modulation of the recovery from desensitization. In contrast, only E413 affected the onset of SSD.

### Mg^2+^ shares binding sites for modulating SSD with Ca^2+^

In a previous study, in which both, Ca^2+^ and Mg^2+^ concentrations were changed at the same time, Mg^2+^ appeared to modulate the ASIC1a pH dependence similarly to Ca^2+ 15^. In our experimental conditions, in the absence of extracellular Ca^2+^, the pH_50_ of activation was 6.59 ± 0.03 (n = 14) at 2 mM Mg^2+^ and 6.72 ± 0.03 (n = 14) at 100 nM Mg^2+^, indicating an alkaline Δ1pH_50_ of 0.13 ± 0.02 pH units with the lower Mg^2+^ concentration (**Fig. 7a**). For the SSD, decreasing the Mg^2+^ concentration from 2 mM to 0.1 mM did not significantly change the pHD_50_. Therefore, the two conditions 10 mM and 0.1 mM Mg^2+^ were compared. A pHD_50_ of 6.97 ± 0.03 (n = 8) was obtained with 10 mM Mg^2+^, while the pHD_50_ at 0.1 mM Mg^2+^ was 7.37 ± 0.03 (n = 8), indicating a shift toward more alkaline pH_50_ values by 0.39 ± 0.01 pH units with the lower Mg^2+^ concentration.

**Figure 7.**
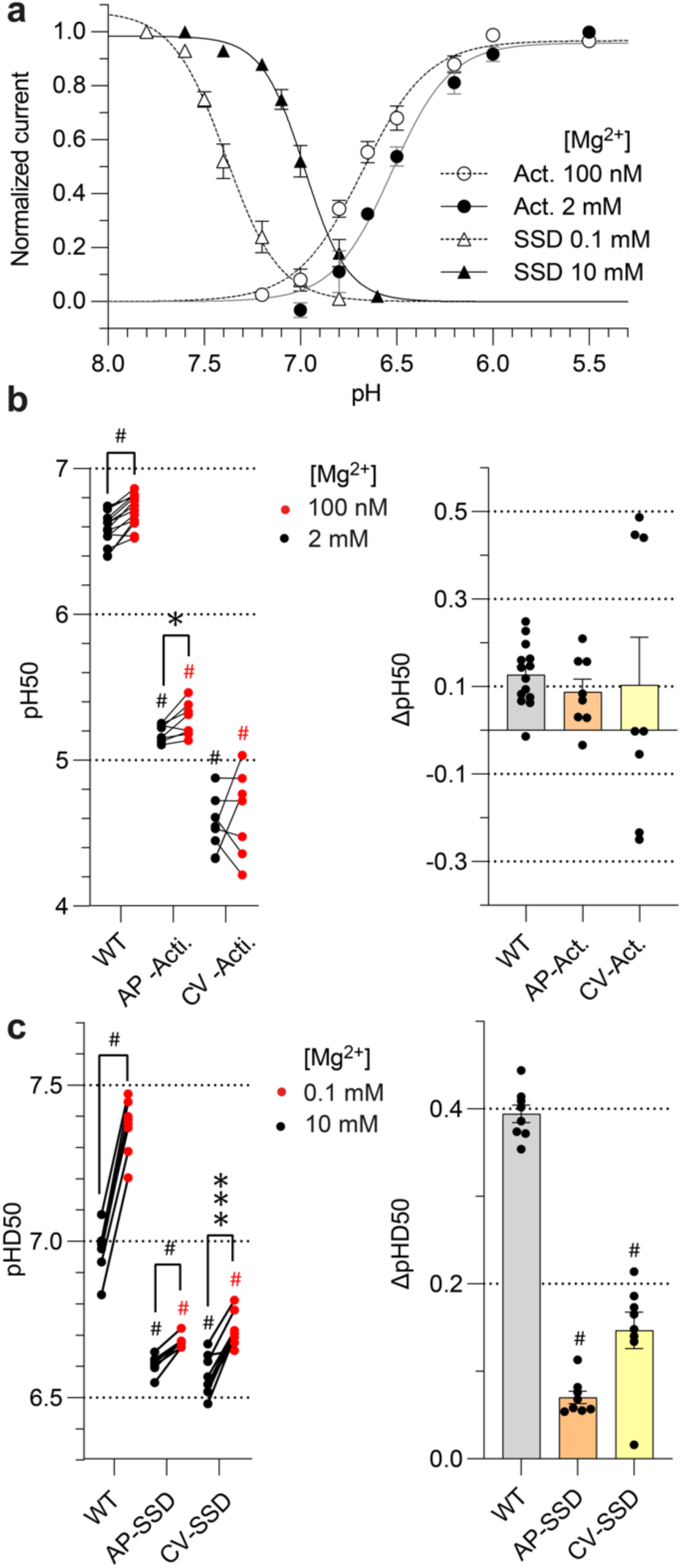
Magnesium modulates ASIC1a activation and desensitization of ASIC1a and shares binding sites with Ca^2+^ for desensitization. **a** pH dependence curves of activation and SSD of ASIC1a (n = 8-9). Activation curves were measured at 100 nM free Mg^2+^ (open circles) or 2 mM Mg^2+^ (filled circles) in the stimulation solution in the absence of Ca^2+^. SSD curves were measured at 0.1 mM Mg^2+^ (open triangles) or 10 mM Mg^2+^ (filled triangles) in the conditioning solution, in the absence of Ca^2+^. The conditioning solution of activation experiments, and the stimulation solution of SSD experiments contained 2 mM Mg^2+^. For the measurement of the activation pH dependence, the pH of the conditioning solution was 7.4, and for the measurement of the SSD pH dependence, the pH of the stimulation was pH5.0. Currents are normalized to the maximum peak current. The lines represent fits to the Hill equation. A representative data set is shown. **b-c** left panels, pH_50_ (**b**) and pHD_50_ values **c** obtained from activation and SSD curves with stimulation **b** or conditioning solutions (**c**) containing low free Mg^2+^ (red symbols) or high free Mg^2+^ (black symbols) as indicated, n = 8. The pH dependence at low and high free Mg^2+^ was measured in the same oocytes. The comparison of the two Mg^2+^ conditions was analyzed with a paired t-test, and the difference to WT values was analyzed with one-way ANOVA followed by Tukey’s multiple comparison test. Right panels, ΔpH_50_ (pH_50,100nM_-pH_50,2mM_, **b**) or ΔpHD_50_ (pHD_50,0.1mM_-pHD_50,2mM,_ **c**) are shown as mean ± SEM, n = 8. Comparison of the mutants to the WT was done by one-way ANOVA and Dunnett’s multiple comparison test. *, p < 0.05; **, p < 0.01; ***, p < 0.001; #, p < 0.0001.

To determine whether Ca^2+^ and Mg^2+^ ions may share binding sites for modulation of the pH dependence, the ΔpH_50_ induced by a change in Mg^2+^ concentration was measured with the previously established combined mutants for Ca^2+^, AP-Act. and CV-Act. (**Fig. 3**). At 2 mM Mg^2+^, the pH_50_ of the two combined mutants was strongly shifted towards acidic values compared to the WT (**Fig. 7b**, left panel), as observed previously in the 2 mM Ca^2+^ conditions (**Fig. 3b**). The ΔpH_50_ between the two Mg^2+^ concentrations obtained with the mutants was not significantly different from the WT value (**Fig. 7b**, right panel), as opposed to the effects of changing the Ca^2+^ concentration on the AP-Act. mutant (**Fig. 3d**). The absence of an effect with the AP-Act. mutant may be due to the fact that in the WT the Mg^2+^-induced pH50 shift was very small. An analogous analysis was carried out for the SSD with the previously described combined mutants AP-SSD and CV-SSD (**Fig. 4**). The pHD_50_ values of the mutants at 10 mM Mg^2+^ were also shifted to more acidic values relative to WT (**Fig. 7c**), as observed with 2 mM Ca^2+^ (**Fig. 4b**). In both mutants, the Mg^2+^-dependent shift was strongly and significantly reduced to 0.07 ± 0.01 (n = 8, AP-SSD) and 0.15 ± 0.02 (n = 8, CV-SSD, **Fig. 7c**). This indicates that Mg^2+^ modulates desensitization and shares binding sites with Ca^2+^ for desensitization.

### Conservation of Ca^2+^ binding sites in ASIC1b

ASIC1b is expressed in the peripheral but not the central nervous system and has a lower apparent affinity for H^+^ compared to ASIC1a ^15, 32^. ASIC1a and ASIC1b are splice variants differing in the N-terminal ∼180 residues, which correspond to the first transmembrane segment, the finger, and parts of the palm and ý-ball domains. The alignment between ASIC1a and ASIC1b (**Supplementary Fig. 2**) indicates that all ASIC1a residues involved in Ca^2+^ regulation, except for Glu97, are conserved in ASIC1b. Therefore, the acidic pocket and pore-lining parts are mostly conserved between ASIC1a and ASIC1b, while parts of the central vestibule are different. Ca^2+^ was shown to shift the SSD curve of ASIC1b similarly to ASIC1a and the activation curves to a lesser extent ^15^. Our measurements confirmed the previous data on ASIC1b, with pH_50_ values of 6.13 ± 0.02 (n = 9) at 2 mM Ca^2+^ and 6.33 ± 0.02 (n = 9) at 100 nM Ca^2+^ (**Fig. 8a**), and pHD_50_ values of 7.13 ± 0.02 (n = 9) at 2 mM Ca^2+^ and 7.37 ± 0.01 (n = 9) at 0.1 mM Ca^2+^. The ΔpH_50_ and ΔpHD_50_ induced by changes in extracellular Ca^2+^ concentration are therefore similar between ASIC1a and ASIC1b. To determine whether the Ca^2+^ binding sites are functionally conserved in ASIC1, the homologous combined mutations to AP-Act., CV-Act., AP-SSD and CV-SSD, previously constructed in ASIC1a, were generated in ASIC1b, with the difference that the non-conserved ASIC1a-E97A mutation could not be included in ASIC1b. The ASIC1b AP-Act. and CV-Act. mutants appeared to have a strong acidic shift of their pH dependence with current amplitudes still increasing at pH3, and a high variability, precluding therefore a precise measurement of the activation pH dependence. For the SSD, the WT ΔpHD_50_ was 0.24 ± 0.02 (n = 9, **Fig. 8b**). The ΔpHD_50_ of the mutant AP-SSD was not different from the WT value (0.17 ± 0.08 (n = 12)), while in CV-SSD, the regulation of the SSD pH dependence by Ca^2+^ was suppressed, with a pHD_50_ value of -0.14 ± 0.09 (n = 10; **Fig. 8b**). These two combined mutants produced transient currents with a sustained current component (**Fig. 8c**). To conclude, the Ca^2+^ binding site for desensitization of the central vestibule is conserved between ASIC1a and ASIC1b. Due to the strong changes in basic properties of the combined mutants for activation, it could not be tested whether the Ca^2+^ binding sites for activation are conserved between the two splice variants.

**Figure 8.**
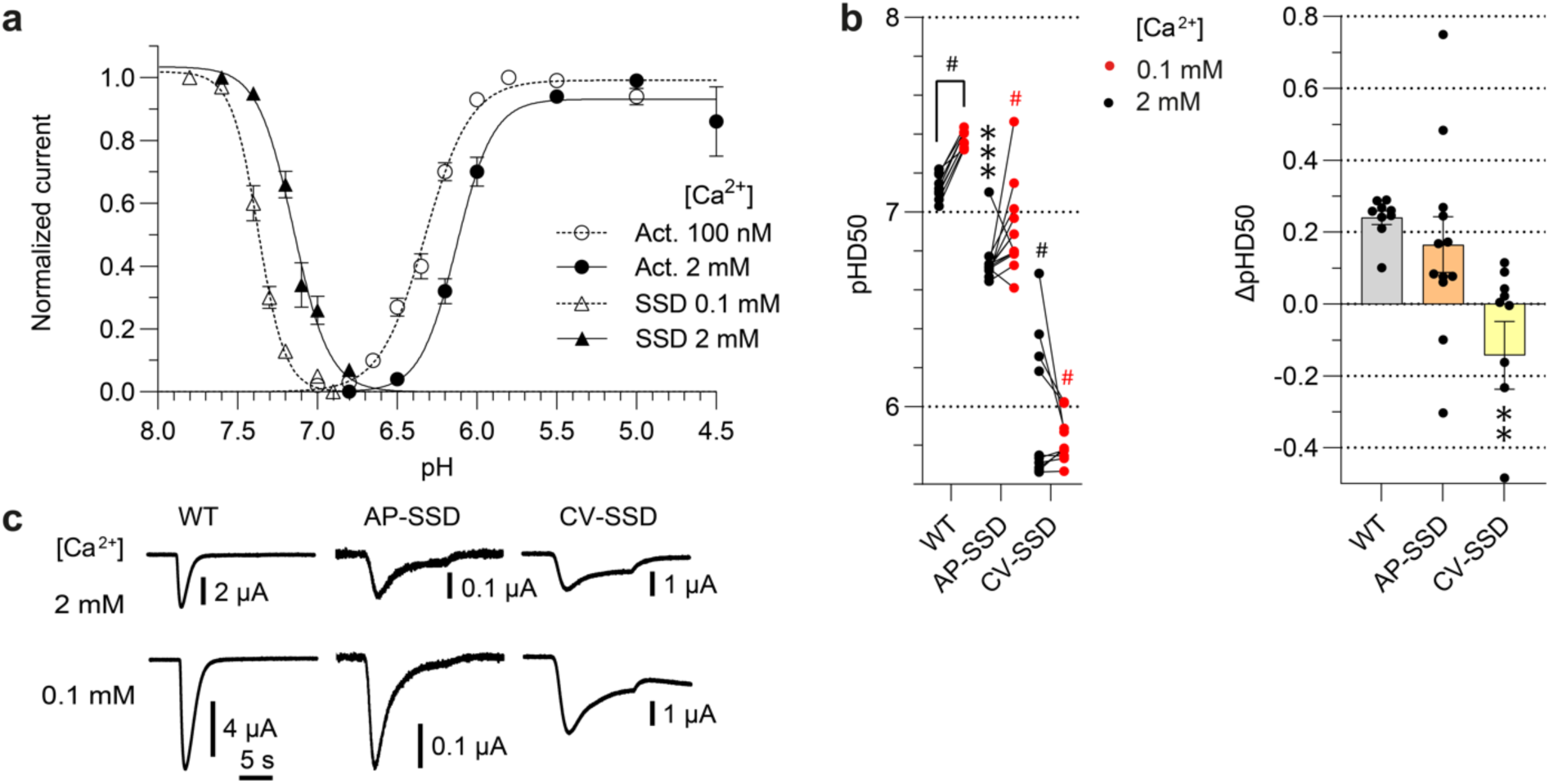
ASIC1b shares Ca^2+^ binding sites for desensitization with ASIC1a in the central vestibule. **a** pH dependence curves of activation and SSD of ASIC1b (n = 9). Activation curves were measured at 100 nM free Ca^2+^ (open circles) or 2 mM Ca^2+^ (filled circles) in the stimulation solution. SSD curves were measured at 0.1 mM Ca^2+^ (open triangles) or 2 mM Ca^2+^ (filled triangles) in the conditioning solution. The conditioning solution of activation experiments and the stimulation solution of SSD experiments contained 2 mM Ca^2+^. Currents were normalized to the maximum peak current. The lines represent fits to the Hill equation. **b** Left panel, pHD_50_ values obtained from SSD curves with conditioning solutions containing 0.1 mM Ca^2+^ (red symbols) or 2 mM Ca^2+^ (black symbols) as indicated, n = 9-12. The comparison of the two Ca^2+^ conditions was analyzed with a paired t-test, and the difference between mutant and WT pHD_50_ values was analyzed with one-way ANOVA followed by Tukey’s multiple comparison test. Right panel, ΔpHD_50_ (pHD_50,0.1mM_-pHD_50,2mM_) values, shown as mean ± SEM, n = 9-12. Comparison of the mutants to the WT was done by one-way ANOVA and Dunnett’s multiple comparison test. *, p<0.05; **, p<0.01; ***, p<0.001; #, p<0.0001. The pH dependence at 0.1 mM and 2 mM Ca^2+^ was measured in the same oocytes. AP-SSD combines the mutations E206A, E225A, Q257A, E229A, D332A and D394A. CV-SSD combines the mutations E360A and E398A. **c** Representative current traces of WT and combined mutants. The traces were obtained at the two Ca^2+^ concentrations, at a pH close to the pH_50_.

## Discussion

Extracellular Ca^2+^ competes with protons for binding sites on ASIC1a and modulates H^+^-induced activation. Based on published crystal structures obtained in the presence of divalent cations and our MD simulations with a structural ASIC1a model, we predicted specific residues of the acidic pocket and the central vestibule to interact with Ca^2+^ ions. Mutation of candidate residues and functional analysis identified several residues of the acidic pocket, the central vestibule and the pore entry as contributing to the modulatory effect of Ca^2+^ (**Fig. 9a-b**), most likely by being part of binding sites. We show here also that Mg^2+^ ions share binding sites with Ca^2+^ for desensitization in both the acidic pocket and the central vestibule. In addition, the Ca^2+^ binding sites for desensitization in the central vestibule are functionally conserved between the splice variants ASIC1a and ASIC1b.

**Figure 9.**
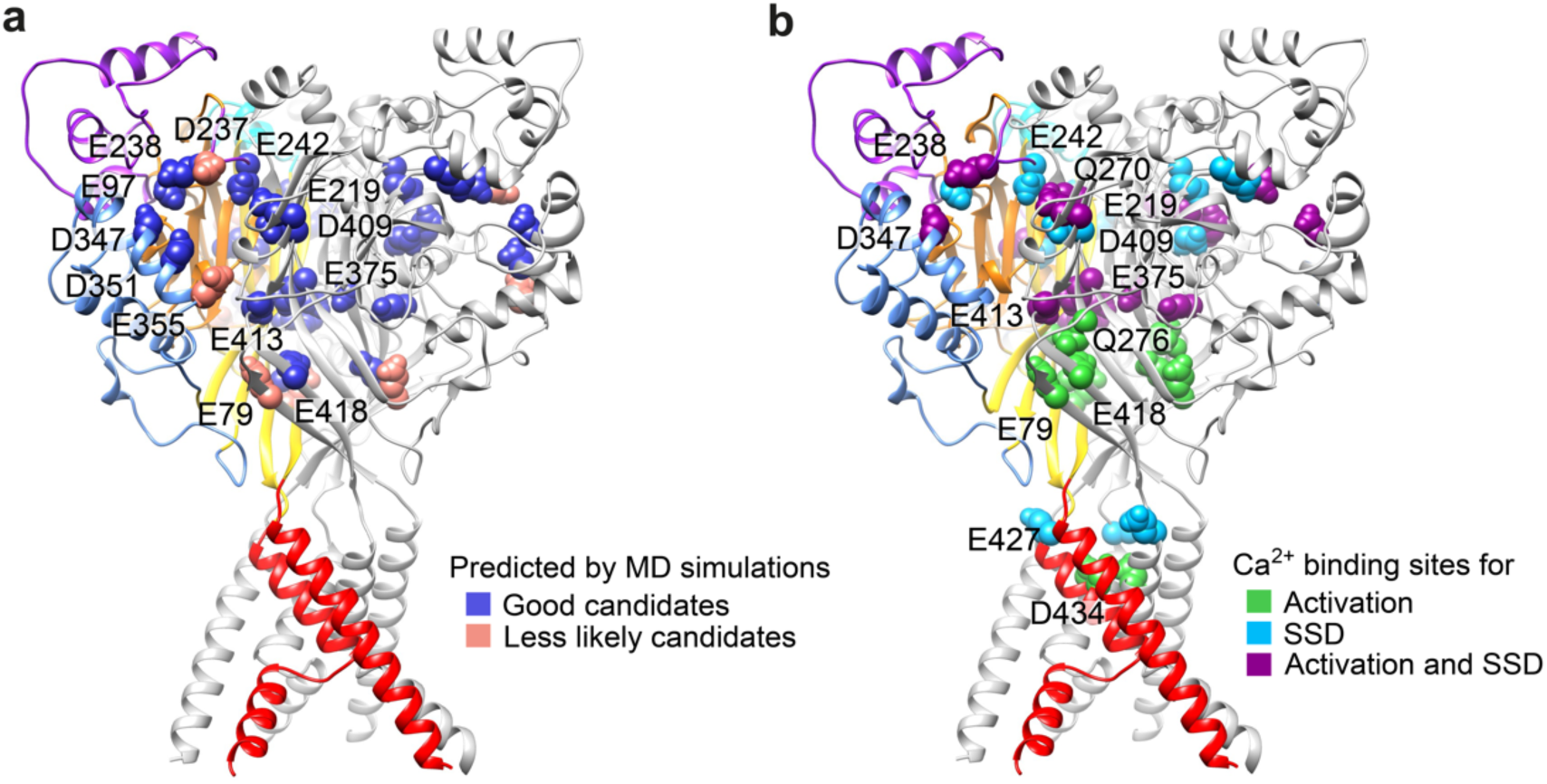
Predicted and confirmed residues for Ca^2+^ binding sites in ASIC1a. Structural images of an ASIC1a trimer in the closed state, from a model based on the crystal structure (5WKU). The different domains are indicated by specific colors in one of the three ASIC subunits, while the two others are represented in grey. **a** Residues predicted by MD simulations as Ca^2+^-binding sites are indicated by two different colors to distinguish the good candidates from less likely candidates. **b** Residues identified by functional analysis as Ca^2+^-binding sites are colored depending on their involvement in activation, SSD or both.

In the mammalian brain, basal interstitial Ca^2+^ and Mg^2+^ concentrations are between 1 and 2 mM. Strong neuronal activity, a seizure or an ischemic stroke can lower the extracellular Ca^2+^ concentration down to 0.1mM ^7, 8, 9^. During ischemia, the anaerobic metabolism produces lactate, which chelates Ca^2+ 33^. Mg^2+^ is one of the most abundant ions and is a cofactor of many enzymes in neurons and glia. Low levels of Mg^2+^ have been associated with pathological conditions such as ischemic stroke, Alzheimer’s disease or migraine headaches ^34^. Also, Mg^2+^ depletion has been reported after brain injury, while Mg^2+^ administration has shown neuroprotective effects ^35^.

Calcium interacts in binding sites primarily with charged amino acid side chains, but main chain carbonyl oxygen atoms can also contribute to coordination ^36^. Even if hydrated, Ca^2+^ can bind with reasonable affinity. In divalent cation binding sites, the coordination structure of Mg^2+^ is generally different from that of Ca^2+^. The ionic radius of Ca^2+^ is with 0.99 Å larger than the one of Mg^2+^ (0.66 Å). Ca^2+^ has therefore a lower charge density with a tendency to coordinate more interacting atoms in binding sites. Mg^2+^ has a higher hydration energy than Ca^2+ 10^, and in binding sites, the distance between the divalent cation and the interacting oxygen atoms is smaller for Mg^2+^ with 2.0 to 2.1 Å than for Ca^2+^ with 2.3 to 2.6 Å ^37^. In troponin C for example, Ca^2+^ has been shown to coordinate an additional glutamate compared to Mg^2+^, which may be critical for the selectivity between these two ions and may explain the better affinity for Ca^2+ 38^. The Ca^2+^-permeable N-methyl-D-aspartic acid glutamate channels are inhibited by physiological Mg^2+^ concentrations by open channel block ^39^, showing also different functions of these two ions. The modulation of ASIC1a by Ca^2+^ and Mg^2+^ is qualitatively similar. We show for ASIC1a that Mg^2+^ shifts the pH dependence less than Ca^2+^ and that Mg^2+^ appears to share binding sites with Ca^2+^ only for desensitization.

Lowering of the extracellular Ca^2+^ concentration increases the activity of voltage-gated Na^+^ channels ^10^, the Na^+^ leak channel (NALCN) ^11^ and the Ca^2+^ homeostasis modulator 1 (CALHM1)^12^. For voltage-gated Na^+^ channels, it was concluded that the effects of Ca^2+^ were due to surface charge screening. The inhibition of NALCN is likely due to pore block ^11^, while Ca^2+^ was shown to interact with an Asp residue of CALHM1 located outside the permeation pathway ^12^.

On ASIC3, lowering the extracellular Ca^2+^ concentration induces a considerably stronger shift of the activation pH dependence than in ASIC1a ^18^. Removal of the extracellular Ca^2+^ at physiological pH7.4 activates ASIC3 ^16^ but not ASIC1a. An ASIC3 gating model was proposed in which acidification opens ASIC3 by inducing unbinding of Ca^2+^ ions from the pore entry ^16^. Consistent with this hypothesis, ASIC3-Glu435, a pore residue not conserved in ASIC1a (the residue at the homologous position in ASIC1a is Gly430), was identified as important for the ASIC3 modulation by Ca^2+ 19^. Recently, our group identified additional residues that contribute to a similar extent as Glu435 to Ca^2+^ modulation of ASIC3 activation, Asp439 (corresponding to Asp434 in hASIC1a) in the pore entry, and Glu212 (corresponding to Glu219 in hASIC1a) of the acidic pocket, showing for the first time a contribution of residues outside the pore entry to ASIC3 modulation by Ca^2+ 18^. Three residues were involved in the Ca^2+^ regulation of ASIC3 SSD, Glu212 and Glu235 of the acidic pocket (corresponding to Glu219 and Glu242 in hASIC1a) and Asn421 of the lower palm (corresponding to Asn416 in hASIC1a) ^18^. Functional observations indicate that ASIC1a is activated by H^+^-induced allosteric changes ^40^. Further support for an allosteric activation mechanism came from voltage-clamp fluorometry experiments which showed evidence for conformational changes linked to ASIC1a activation ^27, 28, 29, 30^ and from the observation that mutation of the two residues involved in ASIC1a pore block by Ca^2+^ did not abolish pH-dependent gating ^17^. The E425G mutation decreased the Ca^2+^-induced shift of the pH dependence of SSD, while D432C decreased the Ca^2+^-induced shift of the activation pH dependence ^17^. The statistical significance of this difference relative to WT was however not analyzed. By mutating the corresponding residues in human ASIC1a, E427 and D434, to Ala, we confirmed these earlier findings, showing that these two residues in the pore are involved in the modulation effect of Ca^2+^.

Compared to ASIC3 in which binding sites in the pore and the acidic pocket contribute to Ca^2+^ modulation of activation, ASIC1a has additional binding sites in the central vestibule involved in activation (**Fig. 9b**). Regarding SSD modulation, the acidic pocket and central vestibule are involved in both ASIC1a and ASIC3, with an additional small contribution of the pore residue Glu427 in ASIC1a (**Fig. 9b**). The comparable effect of Ca^2+^ on ASIC1a and ASIC1b ^15^ suggested conserved binding sites. However, our functional analysis of ASIC1b confirmed only the conservation of Ca^2+^ binding sites in the central vestibule for desensitization. The pH dependence of the combined mutants for activation was too strongly shifted for reliable measurement and the SSD Ca^2+^ modulation of the combined acidic pocket mutant for desensitization was not significantly different from WT.

X-ray crystallography analysis showed that in the high pH resting state of chicken ASIC1a two Ba^2+^ ions bind to each acidic pocket and three to the central vestibule ^20^. In the desensitized state, only one Ba^2+^ site per acidic pocket remained, and no Ba^2+^ sites were found in the central vestibule. It also needs to be considered that ASIC1a opening and desensitization require an acidic pH, which likely leads to protonation and therefore neutralization of the negative charge of some acidic residues in the proximity of Ca^2+^ binding sites, disfavoring cation binding. The Ba^2+^ binding was not determined on the open ASIC1a structure. Since the conformation of the acidic pocket is similar in the open and desensitized state, it is reasonable to assume that each acidic pocket of an open ASIC1a can also accommodate one Ca^2+^ ion. Our MD simulations were carried out with a model of the closed conformation of ASIC1a, with a protonation status mimicking pH7.4. In this conformation, one of the two Ca^2+^ ions of each acidic pocket remained close to E219, D409 and, at a somewhat higher distance, to E242 (**Figs. 2c** and **9a**). We named this position the “inner AP Ca^2+^ site”. The other Ca^2+^ ion had more dynamic interactions, changing its proximity from E97 to mostly D347 and D351, precluding a precise localization of the second Ca^2+^ binding site in the acidic pocket, named here “outer AP Ca^2+^ site”. The single Ba^2+^ remaining in each acidic pocket in the desensitized conformation is located between residues corresponding to D237 and E219 of hASIC1a ^20^, thus overlapping with the inner AP Ca^2+^ site (**Fig. 2c**). The outer AP Ca^2+^ site is therefore only occupied in the resting state, conferring stabilization of the resting state by Ca^2+^.

In the central vestibule, E375 and E413 are located higher up than E79 and E418 (**Fig. 2d**). In many MD trajectories, the Ca^2+^ ions of the central vestibule were closer to E375 and E413 than to E79 and E418. The central vestibule gets narrower during both the closed-open and the open-desensitized transitions ^41, 42^. Therefore, although no Ba^2+^ binding in the central vestibule was observed in the desensitized structure, it can not be excluded that the open ASIC1a could accommodate divalent cations in the central vestibule.

E427 and D434 contribute to the Ca^2+^ binding site in the pore. The conformation of the pore strongly changes upon channel opening. In the closed conformation, the distances between side chain oxygen atoms of different subunits are ∼4 Å for D434 and ∼16 Å for E427, and the distance between the side chain oxygen of D434 and E427 of the same subunit is ∼12 Å. In the open conformation, these intersubunit distances are ∼13 Å for D434 and ∼24 Å for E427. The D434 intersubunit distance of the side chain oxygens in the closed state is with ∼4 Å compatible with Ca^2+^ binding ^43, 44^. However, no divalent binding to the pore was observed in the study by Yoder and Gouaux ^20^ nor by our MD simulation analysis.

The structure and MD simulations had correctly predicted several residues in the acidic pocket and central vestibule contributing to modulation of Ca^2+^ modulation of activation and SSD. The functional analysis showed in addition that the outer AP Ca^2+^ site was more involved in activation (E238, D347), while residues of both acidic pocket sites contributed to Ca^2+^ modulation of SSD (**Fig. 9b**). Regarding the residues of the central vestibule, mutation of all 4 residues impaired Ca^2+^ modulation of activation, with the E79 and in part the E418 mutation exhibiting more pronounced effects compared to mutation of the other two residues. In contrast, only E375 and E413 contributed to Ca^2+^ modulation of SSD. Some residues that appeared to contribute to Ca^2+^ binding sites based on the structural information or MD simulations turned out not to participate in the functional regulation, as D351 and D409 for activation. Despite the close distance of residue D409 from divalent cations in X-ray crystallography analyses and MD simulations, its mutation to Ala even favored Ca^2+^ modulation of activation, as evidenced by the increased shift in the pH dependence of activation. In contrast, the D409A mutation decreased the Ca^2+^-induced shift in the SSD pH dependence. A difference between the structure analyses and the functional studies is the fact that Ba^2+^ was used in the structure determination, which has larger ionic radii than Ca^2+^. The MD simulations were done with the closed conformation of ASIC1a and did therefore not take into account the transitions from the closed to the open or desensitized state, which obviously affected the functional effects of Ca^2+^, as shown by the different effects of mutations on the modulation of activation versus SSD.

The protonation sites governing ASIC1a activation and desensitization are likely located in the acidic pocket, the central vestibule and the wrist/pore entry ^21, 22, 23, 24, 25, 27^. While H^+^ binding promotes the activation and desensitization transitions, Ca^2+^ ions stabilize the closed state, competing thereby with H^+^. Ca^2+^ appears not to affect the transitions or the equilibrium between the open and desensitized states, since there were no obvious differences in desensitization kinetics or sustained current amplitudes between the two tested Ca^2+^ concentrations (traces in **Fig. 3e**). Mutation of several amino acid residues identified as Ca^2+^-binding sites reduced the H^+^-sensitivity at physiological Ca^2+^ concentration, suggesting that they are involved in H^+^-sensing in ASIC1a. Although the combination of the best candidates for Ca^2+^ coordination showed a strong and significant decrease in the Ca^2+^ modulation effect, it did not completely abolish Ca^2+^ modulation. We can therefore not exclude the existence of other Ca^2+^ binding sites. Some combined mutants strongly shifted the H^+^-sensitivity and/or disrupted desensitization. Since the properties of these combined mutants were completely different from those of the WT, they could not be used for a reliable analysis of the effect of the combination of those particular mutations on Ca^2+^ modulation of a normal, transient ASIC current.

In conclusion, a change of Ca^2+^ or Mg^2+^ concentrations occurs in certain physiological or pathological conditions, affecting ASIC activity and thus neuronal signaling. We show here that residues in the acidic pocket, the central vestibule and the pore entry contribute to Ca^2+^ modulation of the ASIC1a pH dependence, which is likely due to competition of Ca^2+^ for protonation sites controlling activation and desensitization.

## Methods

### Molecular Dynamics simulations

The homology models for the molecular dynamics simulations were constructed based on the cryo-EM structure with the accession code 6VTE ^45, 46^, with the termini acetylated and methylated, using in-house CHARMM and Python scripts. The all-atom MD simulations were performed with the CHARMM36 force field, using the GROMACS package version 2020.4. The models were inserted in POPC (1-palmitoyl-2-oleoyl-sn-glycero-3-phosphocholine) bilayers comprising 592 lipids and hydrated (model TIP3^47^) at 150 mMol NaCl. The initial positions of the Ca^2+^ were set to correspond to the positions published in the study by Yoder et al. ^20^. Two such systems were combined in an antiparallel way to form a double bilayer system, simulating a cell membrane separating two different water compartments ^48^, resulting in a box containing two channels, 1184 lipids and a total of approximately 690’000 atoms. Before proceeding with the main trajectories, the pKas of all titratable residues were calculated as done previously ^49^. Briefly, short 10 ns unrestrained simulations were produced using the CHARMM 36 force field, and frames were extracted at 2 ns intervals. The duration of these simulations was chosen to ensure adequate relaxation of ions, water molecules, and partial side-chain movements, while avoiding significant conformational changes of larger molecules. For each frame that was extracted, the PBEQ solver implemented in CHARMM ^50^ was utilized to calculate the pKas. The calculated average pKas were then used to determine the initial protonation states of titratable residues to mimic a pH environment of 7.4. After setting the residue protonation states, the system was minimized and equilibrated in six steps with decreasing restrains of the heavy atoms, for a total time of 1.5 ns. A first unrestrained simulation was then conducted for 100 ns. The structure obtained at 100 ns was then used for a second pKa calculation, aimed at identifying residues requiring a modification of their protonation state because of conformational changes of displacements of ions. After updating the residue protonation states, the system was again minimized and equilibrated in six steps with decreasing restraints on the heavy atoms, for a very short total time of 110 ps. Since the structure was extracted from an equilibrated MD simulation, the aim here was solely to allow for the surroundings of the modified sidechains to relax slightly before the simulation. This procedure was repeated five times until the completion of 600 ns of simulation.

### Multi-step computational determination of candidate residues interacting with the Ca^2+^ ions

Since classical MD represents the electronic structure of atoms as spheres, our analysis did not aim to characterize the calcium electronic coordination *sensu stricto.* Coordination numbers or chelation modus were not investigated. Our project consisted of identifying negatively charged residues harboring the most consistent interactions with the divalent ion, inferred through comparison of duration of contact between the center-of-mass of the carbonyl groups and the ion. We hypothesized that during a simulation, it is theoretically possible that a Ca^2+^ ion coordinated by a given ensemble of residues, might leave this ensemble and get captured by another one. To avoid that such a new interaction being ignored, we conducted a two-step blind research of interacting pairs of Ca^2+^ ion – acidic residue.

In the first step, the distances between all Ca^2+^ ions and all carbonyl groups were extracted at 10 ns intervals during each individual 100 ns long simulation. Since two proteins were simulated in the same box, this accounts for 2 (two proteins) × 18 (Ca^2+^) × 183 (Glu and Asp) × 6 (individual 100 ns long simulations) × 10 (ten measurements during 100 ns) = 395280 distances. For the second step, any pair harboring at least one occurrence with a distance Ca^2+^-carbonyl group smaller than 10 Å was retained. These ∼ 600 selected pairs were then subjected to the same distance calculation, but at intervals of 400 ps for a better precision. To further isolate candidate residues, the distance threshold was reduced to 6 Å and residues were retained only if this distance criteria was met during at least 10% of the simulated time, i.e., during at least 10ns during a 100 ns long trajectory. This second criteria did not require the interaction to be continuous in time.

### Molecular biology

The human ASIC1a clone (GenBank U78181 ^51^, in which the mutated residue Asp212 had been corrected to Gly ^52^), rat ASIC1b-M3 (Genbank AJ30992; which was transcribed from the third Met, which corresponds to the first Met of ASIC1a ^53^) and derived mutants were subcloned in the pSP65-derived vector pSD5 that contains 5’ and 3’ non-translated sequences of β-globin to improve the protein stability in *Xenopus laevis* oocytes. Mutants were generated by site-directed mutagenesis using the QuikChange approach, with KAPA HiFi HotStart PCR polymerase (KAPA Biosystems). Combined mutants were synthesized by Genscript. Isolation of high-copy plasmid DNA from *E. coli* was done using NucleoSpin Plasmid (MACHEREY-NAGEL). The mutations were verified by sequencing (Microsynth) and transcription was achieved using the mMESSAGE mMACHINE kit (Thermo Fisher Scientific).

### Xenopus laevis oocyte preparation and use

All animal experiments were carried out in accordance with Swiss laws and were approved by the veterinary service of the Canton de Vaud. 1.3 gL^-1^ of MS-222 (Sigma-Aldrich) was used to anaesthetize female *Xenopus laevis* frogs. The oocytes were extracted by a small incision on the abdominal wall and the lobe was treated with 1mg·ml^-1^ collagenase for the isolation and defolliculation. Healthy stage V and VI oocytes were selected. Oocytes were injected with 50nl of cRNA at 5-1100 ng·μL^-1^. Prior to electrophysiological measurements, the oocytes were stored in Modified Barth’s saline containing (in mM) 85 NaCl, 1 KCl, 2.4 NaHCO_3_, 0.33 Ca (NO_3_)_2_, 0.82 MgSO_4_, 0.41 CaCl_2_, 10 HEPES and 4.08 NaOH at 19°C.

It has been reported that lowering of the extracellular Ca^2+^ concentration can induce inward currents in *Xenopus Laevis* oocytes due to activation of endogenous connexin hemichannels^54^. We have recently shown that a switch at pH7.4 from a solution containing 2 mM of Ca^2+^ to a solution containing 100 nM Ca^2+^ induces a slowly developing inward current in non-injected oocytes ^18^. Due to the small amplitude of this current, its effect on the pH_50_ of expressed ASICs was negligible. We have repeated these control experiments with oocyte batches used in the current study (**Supplementary Table 1 and Supplementary Fig. 3**). Decreasing the Ca^2+^ concentration from 2 mM/pH7.4 to 100 nM produced an inward current of -54 ± 8 nA at a test pH6.0 (n = 16; **Supplementary Fig. 3a-b**) and 1 ± 3 nA at test pH7.4 (n = 16). Measurements at different pH conditions at the Ca^2+^ concentrations of 0.1 mM and 100 nM showed inward currents of the order of 100 nA or less (**Supplementary Fig. 3A and Supplementary Table 1**), suggesting that the presence of these low endogenous currents does not affect the pH dependence measurements of ASIC currents that were generally of several μA.

### Electrophysiological measurements

Electrophysiological recordings were performed 1-3 days after cRNA injection. Whole-cell currents were recorded by two-electrode voltage-clamp using two glass electrodes filled with 1 M KCl with a resistance < 0.5 MΩ. A Dagan TEV200 amplifier (Minneapolis, MN), equipped with two bath electrodes, was used together with an InstruTECH LIH 8+8 interface and PatchMaster software (HEKA-Harvard Bioscience) to perform recordings. Current were recorded at a holding potential of -60 mV, at a sampling interval of 20 ms and filtered at 2 kHz. Oocytes were perfused with experimental solutions by gravity at a flow rate of 8–12 mL·min^−1^, using the cFLow 8 channel electro valve unit (CellMicroControls) with an eightfold perfusion head. The oocytes were exposed to a stimulation solution with a selected pH for 5–10 s once per minute and the pH-current curves were fitted to the Hill equation. Measurements of the pH dependence or kinetics were done in paired experiments, comparing two divalent cation concentrations in the same oocyte. The recording solution contained (in mM): 110 NaCl, 10 HEPES for pH ≥ 6.8 and the indicated Ca^2+^ or Mg^2+^ concentration. HEPES was replaced by MES for solutions with a pH < 6.8 and by Glycine for solutions with a pH ≤ 4. NaOH or HCl was used to adjust the pH. Solutions with 100 nM free Ca^2+^ or Mg^2+^ contained 10 mM EDTA for pH > 6.0 and 20 mM EDTA pH ≤ 6. Total Ca^2+^ or Mg^2+^ concentrations were determined based on the MaxChelator program (https://somapp.ucdmc.ucdavis.edu/pharmacology/bers/maxchelator/webmaxc/webmaxcS. htm^55^) to obtain the desired free Ca^2+^ concentration. No chelator was added in solutions with ≥ 0.1 mM of Ca^2+^.

### Statistics and Reproducibility

Graphpad Prism (version 10) was used for the fits and the statistical analyses. For pH dependence, currents were normalized to the maximum peak current induced. Statistical differences were analyzed with one-way ANOVA test followed by a Dunnett’s or Tukey’s multiple comparisons test for comparison of ≥ 3 groups, and a paired t-test for direct comparison of two groups. The statistical tests applied are indicated in the respective figure legends. Each experiment was repeated measuring at least on two different days and in different batches of oocytes from a different frog. Data are presented as individual data points or as mean ± SEM.

## Data and code availability

All experimental data are contained in the manuscript and supplementary files. Raw data are provided in the supplementary file “figure_raw_data.xlsx”. No unique code was written to generate the data of this study.

## Supporting information

Figure raw data

## Acknowledgements

We thank Eleonora Centonze, Marc Bohnet and Mina Hanna for comments on the manuscript. This work was supported by the Swiss National Science Foundation grant 310030_207878 to S.K. This work was supported by computational resource grants from the Swiss National Supercomputing Centre (CSCS) under project IDs s1037 and s1099.

## Author contributions

Conceptualization, O.M., O.B., S.K.; methodology and investigation, O.M., O.B.; supervision: S.K.; writing original draft, O.M., O.B., S.K.; writing, review & editing: O.M., O.B., S.K.

