## Supplementary material for "Identification of the modulatory Ca^2+^ binding sites of acid-sensing ion channel 1a"

| | 100 nM $Ca^{2+}$ | | | 0.1 mM $Ca^{2+}$ | | | |
|---|---|---|---|---|---|---|---|
| **pH** | **Current amplitude (nA)** | | | **Current amplitude (nA)** | | | **n** |
| **8.5** | 7 | ± | 2 | -9 | ± | 5 | 16 |
| **8.0** | 2 | ± | 3 | -13 | ± | 5 | 16 |
| **7.4** | 4 | ± | 3 | -20 | ± | 5 | 16 |
| **6.5** | -126 | ± | 38 | -26 | ± | 7 | 16 |
| **6.0** | -90 | ± | 34 | -18 | ± | 3 | 16 |
| **5.0** | -42 | ± | 14 | -30 | ± | 5 | 16 |

**S1 Table. Absolute current amplitudes of non-injected *Xenopus laevis* oocytes induced by lowering of the extracellular $Ca^{2+}$ concentration.** Current responses to the indicated $Ca^{2+}$ concentrations and pH conditions measured from a conditioning solution of pH7.4 / 2mM $Ca^{2+}$ at a holding voltage of -60mV.  Maximal current amplitudes were measured and are shown as mean ± SEM.

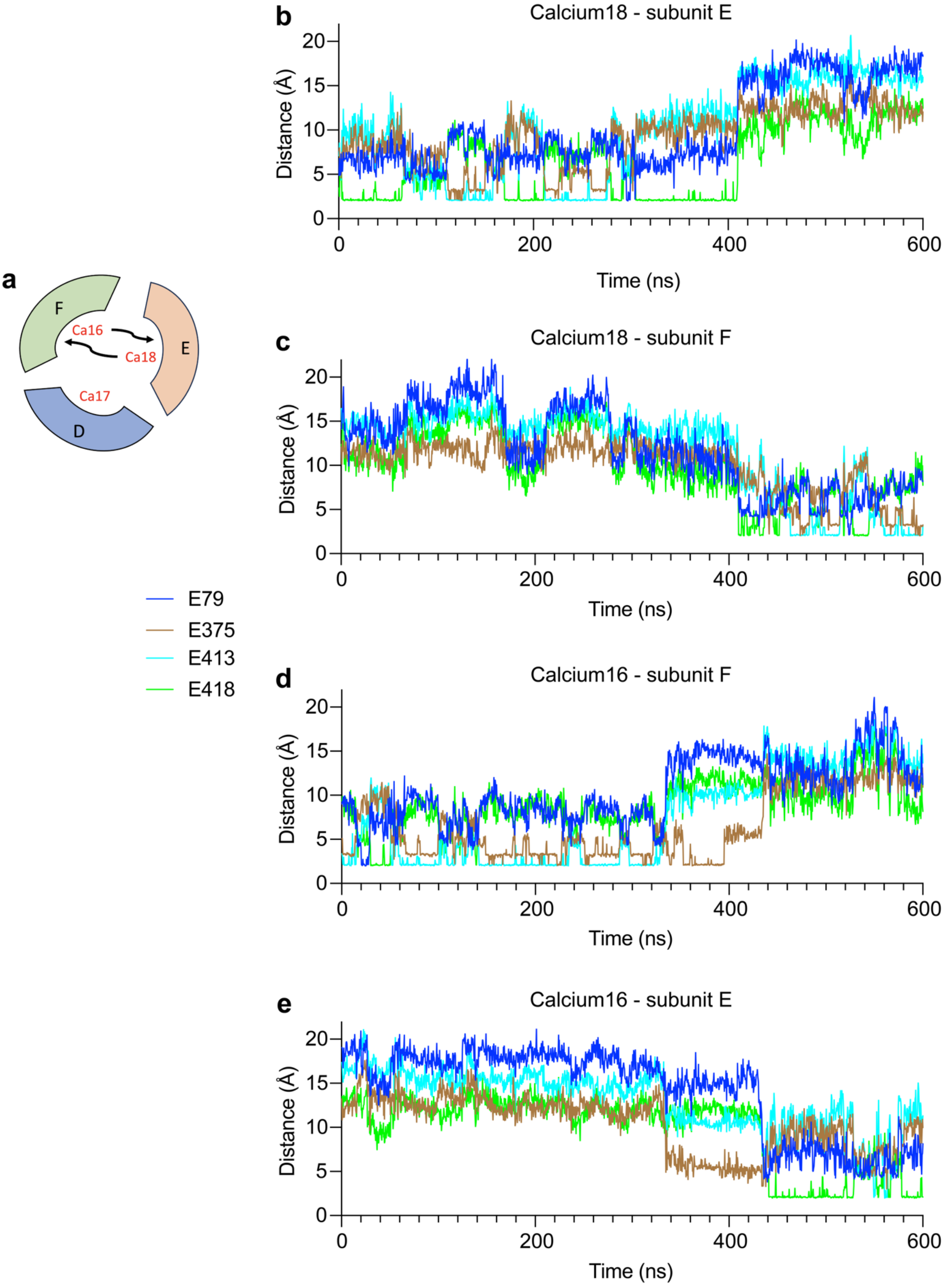


**Supplementary Figure 1. Subunit swapping of $Ca^{2+}$ ions in the central vestibule.** MD simulations were carried out over a total duration of 600 ns (*Materials and Methods*). **a** Illustration of the initial positions of $Ca^{2+}$ ions 16, 17 and 18 in the central vestibule of the channel composed of the subunits D, E and F. The arrows indicate the exchange of positions of Ca16 and Ca18 during the simulation. **b-e** Distances, measured between $Ca^{2+}$ ions and the center-of-mass of the carbonyl groups of acidic side chains of the indicated residues are plotted as a function of simulation time between Ca18 and subunit E (**b**), Ca18 and subunit F (**c**), Ca16 and subunit F (**d**) and Ca16 and subunit E (**e**).

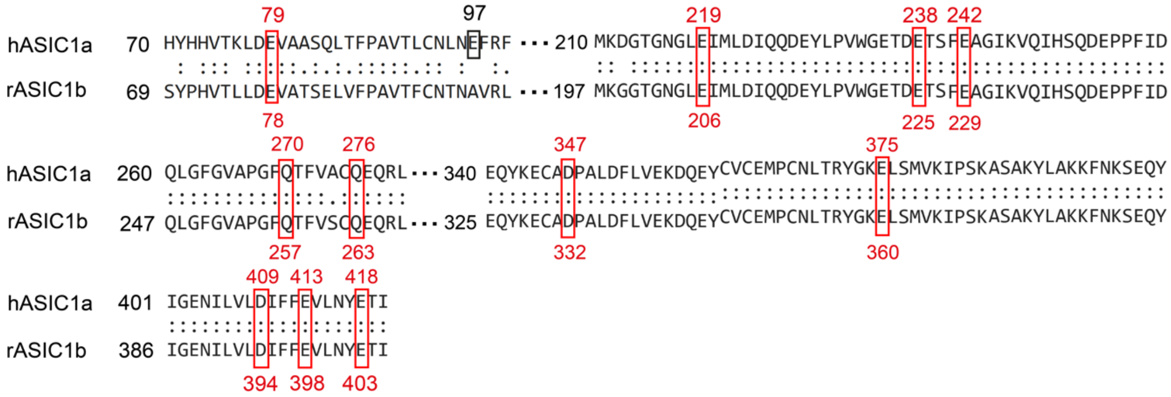


**Supplementary Figure 2. Alignment of amino acid sequences of hASIC1a and rASIC1b in different channel domains.** Corresponding conserved candidate $Ca^{2+}$-binding residues are highlighted by red frames.

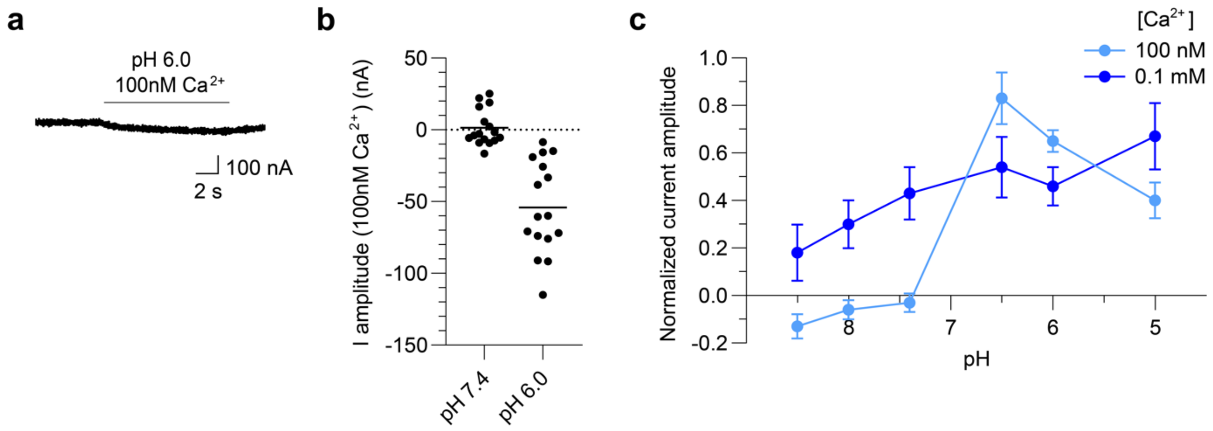


**Supplementary Figure 3. Ionic currents of non-injected *Xenopus laevis* oocytes induced by lowering of the $Ca^{2+}$ concentration. a** Representative current trace from a non-injected *Xenopus* oocyte induced by application of a test pH 6.0 at 100 nM free $Ca^{2+}$, preceded by exposure to a conditioning pH 7.4 at 2 mM $Ca^{2+}$. **b** Current amplitudes were obtained by applying test solutions containing 100 nM free $Ca^{2+}$ at the indicated pH, n = 16. **c** pH dependence curves of activation of endogenous currents measured at 100 nM free $Ca^{2+}$ (light blue filled circles) and 0.1 mM $Ca^{2+}$ (dark blue filled circles) in the test solution from a conditioning solution at pH7.4 containing 2 mM $Ca^{2+}$ (n = 16). In all these experiments, the maximal current amplitude was measured.
